# Tracking influenza a virus infection in the lung from hematological data with machine learning

**DOI:** 10.1101/2022.02.23.481638

**Authors:** Suneet Singh Jhutty, Julia D. Boehme, Andreas Jeron, Julia Volckmar, Kristin Schultz, Jens Schreiber, Klaus Schughart, Kai Zhou, Jan Steinheimer, Horst Stöcker, Sabine Stegemann-Koniszewski, Dunja Bruder, Esteban A. Hernandez-Vargas

**Author notes:** **Contributions**. These authors contributed equally: Suneet Singh Jhutty, Julia D. Boehme. These authors jointly supervised this work: Dunja Bruder, Esteban A. Hernandez-Vargas.

## Abstract

The tracking of pathogen burden and host responses with minimal-invasive methods during respiratory infections is central for monitoring disease development and guiding treatment decisions. Utilizing a standardized murine model of respiratory Influenza A virus (IAV) infection, we developed and tested different supervised machine learning models to predict viral burden and immune response markers, *i*.*e*. cytokines and leukocytes in the lung, from hematological data. We performed independently *in vivo* infection experiments to acquire extensive data for training and testing purposes of the models. We show here that lung viral load, neutrophil counts, cytokines like IFN-γ and IL-6, and other lung infection markers can be predicted from hematological data. Furthermore, feature analysis of the models shows that blood granulocytes and platelets play a crucial role in prediction and are highly involved in the immune response against IAV. The proposed *in silico* tools pave the path towards improved tracking and monitoring of influenza infections and possibly other respiratory infections based on minimal-invasively obtained hematological parameters.

## INTRODUCTION

Respiratory infections by influenza (flu) viruses cause 3 to 5 million cases of severe illness every year^1^. Influenza A virus (IAV) is especially severe among high-risk groups like the elderly, infants, pregnant women, and immunocompromised people^2^. Next to its high prevalence in annual epidemics, IAV has led to high mortality during several pandemics including the Spanish flu in 1918 and more recently the swine flu in 2009^3,4^. Generally, the outcome of flu disease highly depends on viral factors as well as host immunity. Accordingly, a fatal course of infection can result from either insufficient control of viral spread, hyperinflammation, and/or a secondary bacterial infection^5^. Thus, tracking of viral burden as well as host responses in the lungs is important for monitoring IAV pathogenesis and tailoring targeted therapies.

Methodologically, diagnosis and tracking of acute IAV infection can be performed by assessing viral antigen, nucleic acid, or infectious particles from upper or lower airway lavages, aspirates, or swabs. Likewise, monitoring of lower airway immune responses is accomplished by *e*.*g*. quantification of inflammatory cytokines or leukocytes in bronchoalveolar lavage fluid (BALF). Next to several obvious disadvantages of these methods (low sensitivity, costly and time-consuming analyses, and/or high technical requirements), the biggest hurdle lies in the invasive sampling procedure that poses a risk to the acutely infected patient^6^. Accordingly, the development of non-or minimally invasive approaches that allow observation of the disease status during IAV infection remains an unmet medical need.

Besides inducing acute inflammatory responses in the airways, influenza infection results in peripheral immune activation manifesting in altered blood cell composition^7^, transcriptional signatures, and cytokine and chemokine levels in mice and humans^8–10^. While the severity and longitudinal analyses revealed distinct molecular and/or cellular characteristics in peripheral blood of IAV infected hosts, the suitability of each of these markers (and blood parameters in general) for predicting the disease status is still unknown.

Here, we propose for the first time a framework intertwining *in vivo* experiments and machine learning methods to forecast IAV infection parameters in the lung from blood sample data that can be accessed minimal-invasively. To this end, we employed different machine-learning models for blood-lung mapping. Experimentally, we utilized an established mouse model of sublethal respiratory IAV infection^11,12^ and simultaneously assessed the kinetics of pathogen burden, lung inflammation, as well as systemic cellular changes following infection. Ultimately, several independent *in vivo* experiments were used to validate the applicability of the proposed framework. Our primary computational approach was deep learning, which represents a class of machine learning algorithms that uses multiple layers of information processing for feature extraction and pattern analysis^13,14^. These methods have already been successfully applied in several biological fields including the prediction of transcriptional enhancers^15^, protein secondary structure^14^, and the pathogenicity of genetic variants^17^. The image recognition abilities of machine learning algorithms have already been tested for their diagnostic value and show promising results^18,19^.

## RESULTS

Our methodology consisted of three consecutive stages: *in vivo* experimental data acquisition, model training, and independent experimental model validation (Figure 1). For the acquisition of experimental data, mice were infected with a sublethal dose of PR8 (H1N1) followed by the quantification of lung viral load, pulmonary innate and adaptive leukocyte subsets, pulmonary cytokine levels, and hematological parameters over a total period of 11 days. Blood and lung parameters were measured (Figure 1) in two independent experiments, mice (*n* = 4-6 per time point) were sacrificed on days 1, 2, 3, 4, 5, 7, 9, and 11 post-infection (pi). These experimental data built the basis for model training using different machine learning approaches to identify the relationship between hematological and pulmonary parameters and to train and optimize the model accordingly. To validate the predictive value of our model, we performed two additional independent infection experiments with mice sacrificed on days 2, 4, 6, 9, and 11 pi and used the mathematical models to predict the lung viral burden, leukocyte composition, and cytokine levels, respectively, based on experimental hematological parameters.

**Figure 1:**
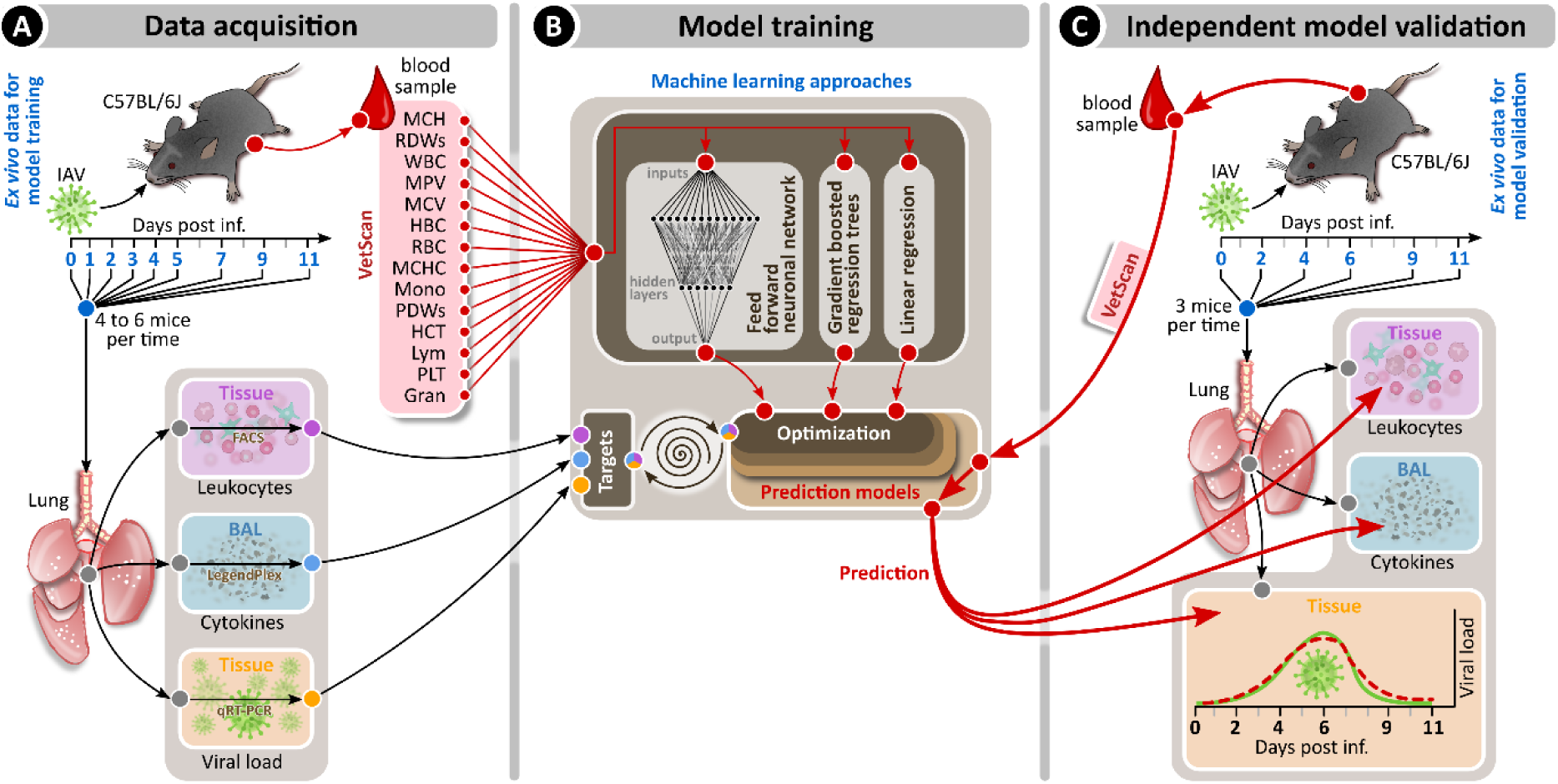
Experimental scheme for the machine learning approaches of the respiratory IAV infection. Mice were intranasally infected with a sublethal dose of IAV PR/8/34 on day 0 and sacrificed on the indicated days. Blood was collected for hematology analyses (Supplementary Figures S1-2), bronchoalveolar lavage was performed to analyze lung cytokines and lung tissue samples were used to monitor either viral load (experiment 1, Supplementary Figure S5) or pulmonary leukocyte subsets (experiment 2, Supplementary Figures S6, S8) (A). The hematological data from this initial set of experiments were used to build and train different machine learning models (B). Data from a separate experiment (Supplementary Figures S3-5, S7, S9) were used for testing and evaluation of machine learning algorithms (C).

### Blood and Lung Data Analysis

In a first step, we conducted a correlation analysis to uncover potential linear relationships between selected hematological and pulmonary parameters (Figure 2 and Supplementary Figures S10-12). We found a very strong correlation (Pearson correlation coefficient > 0.9) between blood leukocytes and lymphocytes, which can be attributed to the fact that lymphocytes constitute the largest leukocyte fraction in mice^20^. Likewise, a strong correlation observed between hematocrit as well as hemoglobin and erythrocyte numbers (Pearson correlation coefficient > 0.8) can be attributed to the fact that hematocrit, as well as hemoglobin, are red blood cell-associated parameters^21^.

**Figure 2.**
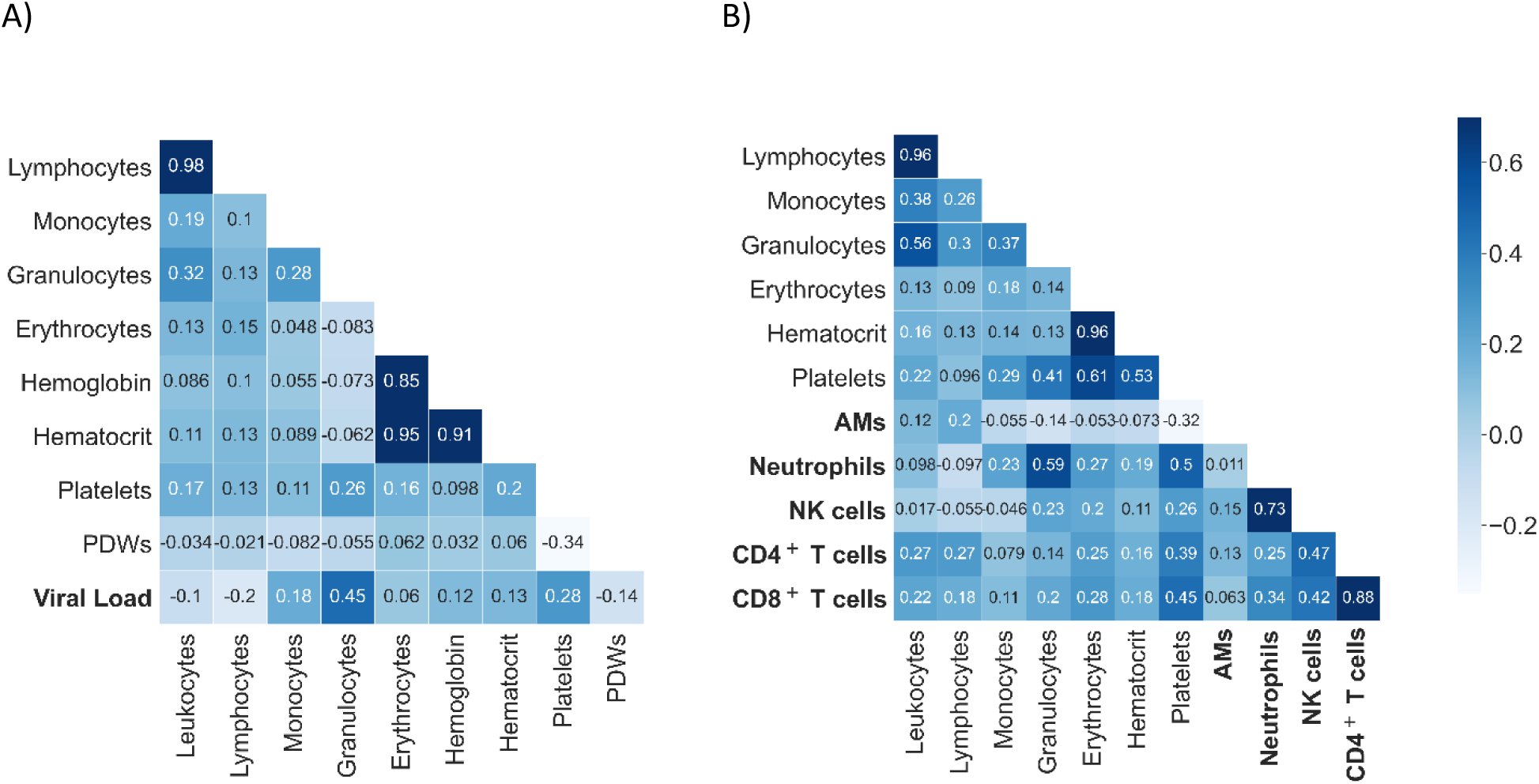
Selected correlations of blood cells, lung leukocytes, and lung viral load for influenza infection. (A) shows the correlation of blood cells with lung viral load. (B) shows the correlation of blood cells with lung leukocytes. The matrices depict the respective Pearson correlation coefficients from the initial experiments used as training data for the machine learning models. We observed some strongly related clusters like erythrocytes, haemoglobin, and hematocrit or NK, CD4^+^ T, and CD8^+^ T cells. IAV-associated lung markers that were later predicted are shown in bold letters. All other parameters were provided to the algorithm to make the estimation. Here, only a small subset of the data and its correlations are shown. To view all the data with its correlations, please refer to the supplemental material (Supplementary Figures S10-12).

Strikingly, we did not observe any strong correlation between blood and lung compartment variables. However, there were strong correlations within cells of the lung compartment, such as CD4^+^ T cells and CD8^+^ T cells as well as neutrophils and natural killer (NK) cells. As a linear correlation does not include higher-order nonlinear temporal relations, we next employed machine learning algorithms and compared the obtained results with the linear regression model.

### Tracking IAV Infection in the Lungs from Blood-Derived Parameters

To predict influenza virus levels and immunological markers in the lungs from hematological parameters, we employed feedforward neural networks, gradient boosted regression trees, and a linear regression model. These algorithms considered 14 hematological parameters (see Table 1) as features to predict the respective target lung variable. The main scores used for comparing and evaluating different machine learning algorithms were based on the average squared difference between the estimated values and the actual value (mean squared error). The proportion of the variance in the dependent variable that is predictable from the independent variables was based on the R^2^ score. Overfitting was reduced with regularization techniques presented in methods.

**Table 1:** Hematological parameters used to estimate the viral load in the lungs of mice. Blood variables are listed in order of their Pearson correlation with viral load (lowest coefficient: MCH, highest coefficient: granulocytes).

| Blood variable | Interpretation |
| --- | --- |
| MCH (mean corpuscular hemoglobin) | average amount of hemoglobin per red blood cell |
| RDWs (red cell distribution width) | degree of variation in size and shape of red blood cells |
| Leukocytes (white blood cells) | protect against infectious diseases and foreign bodies |
| MPV (mean platelet volume) | average size of platelets in blood |
| MCV (mean corpuscular volume) | average volume of red cells |
| Hemoglobin | oxygen carrier in red blood cells |
| Erythrocytes (red blood cells) | oxygen transportation to the tissue |
| MCHC (MCH concentration) | concentration of hemoglobin in red blood cells per volume |
| Monocytes | subtype of white blood cells |
| PDWs (platelet distribution width) | indicates variation in platelet size |
| Hematocrit | volume percentage of red blood cells in blood |
| Lymphocytes | subtype of white blood cells |
| Platelets | blood component that helps stopping bleeding |
| Granulocytes | subtype of white blood cells |

Figure 3A illustrates the best model prediction of viral levels in the lung, *i*.*e*. the feedforward neural network, based on experimental haematological data. We observed that the model had a good qualitative behavior of the lung viral load (measured as copy numbers of NP transcripts) over the experimental time-frame of eleven days. Day 0 represents the control group. The prediction seemed to be most accurate around the peak of viral replication, *i*.*e*. days 4 and 5 pi. Quantitatively, there was a high variation in the predictions. This was attributed to the variation found in our infection experiments (Figure S1-S9), which is a common observation *in vitro* and *in vivo* viral infection experiments^22–27^. While for some animals there was a large difference between the actual experimental data and the respective predictions in the testing experiments, overall, the qualitative performance on the testing set was good (Figure 3B).

**Figure 3.**
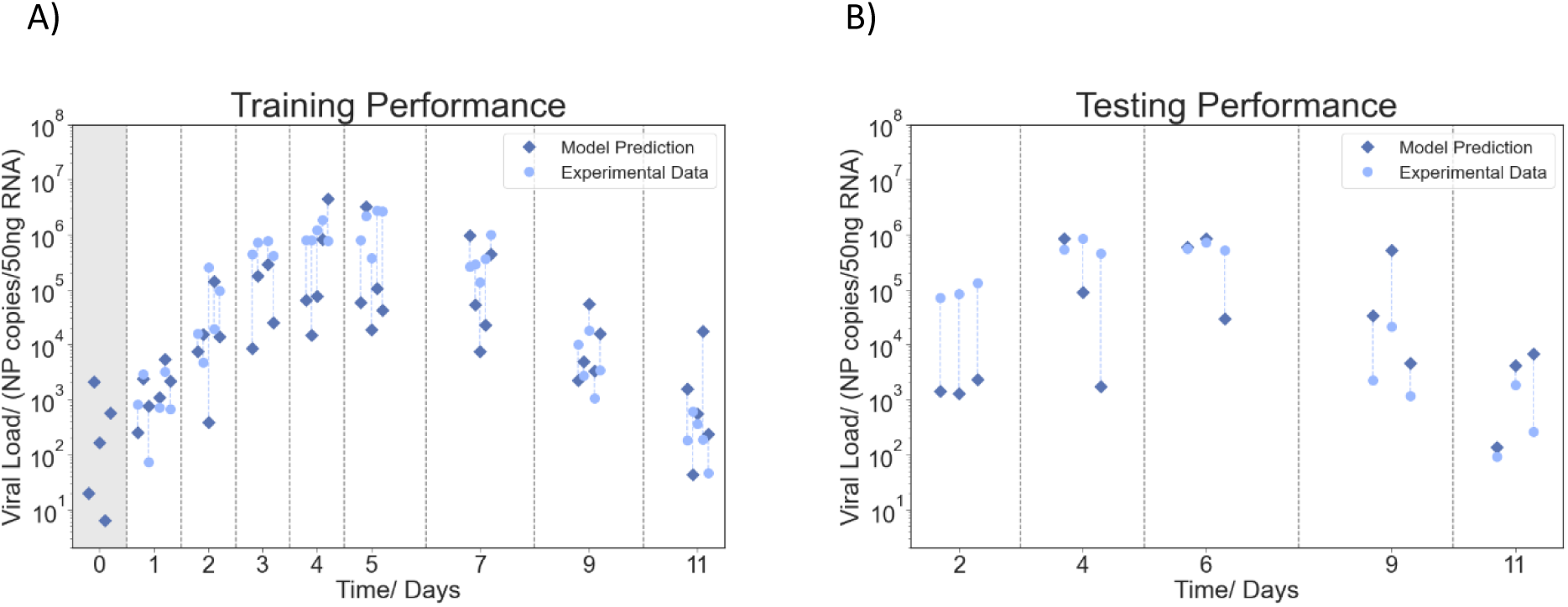
Mapping of the lung viral load from blood data. The plots show the training and testing performance of the neural network model. (A) shows the performance of the model on the training data. (B) shows the performance of the testing data obtained from a second experiment. Each circle represents one mouse, with its matching individual prediction indicated by a connected (blue dashed line) diamond. The vertical lines divide experimental days. Day 0 marks the control group (viral load below measurable threshold) and is highlighted in grey.

In addition to predicting the lung viral load, we tested several machine learning algorithms to predict target lung leukocytes and cytokines from the hematological parameters (Table 2). The comparison in Figure 4A shows how the respective best model performed for different targets (for all results please refer to the supplemental material, Supplementary Figures S13-15). As a benchmark, we used the mean for each target variable calculated from the training data. In almost all cases, the best model performed better than the benchmark. A positive R^2^ score demonstrates the explanatory power of the model. Predictions for lung IFN-γ, viral load, IL-6, and neutrophils were able to outperform the benchmark (Figure 4). Predictions for other lung target parameters are presented in Supplementary Figure S13-S15. Notably, the accuracy of the model predictions was dependent on the stage of the infection. For example, neutrophil numbers in the lungs were better predicted in the later days of infection, while IL-6 and IFN-γ levels were more accurately predicted at the peak of infection (Figure 4B-D). This was also observed for other immune cells such as CD4^+^ and CD8^+^ T cells (Figure S13). Table 2 presents a summary of the best models for predicting the different lung target parameters.

**Table 2.** Best performing model and respective scores for different targets from the lung milieu. Hematological data was used in all cases as input to the algorithms. For each target variable to estimate, different machine learning models were tested.

| Target Variable | Model | MSE | R<sup>2</sup> |
| --- | --- | --- | --- |
| Viral Load | Feedforward NN | 7.53 | 0.18 |
| Neutrophils | LR with PCA | 0.98 | 0.25 |
| IFN-γ | Feedforward NN | 7.18 | 0.15 |
| Interleukin-6 | GBRT | 4.55 | 0.21 |

**Figure 4.**
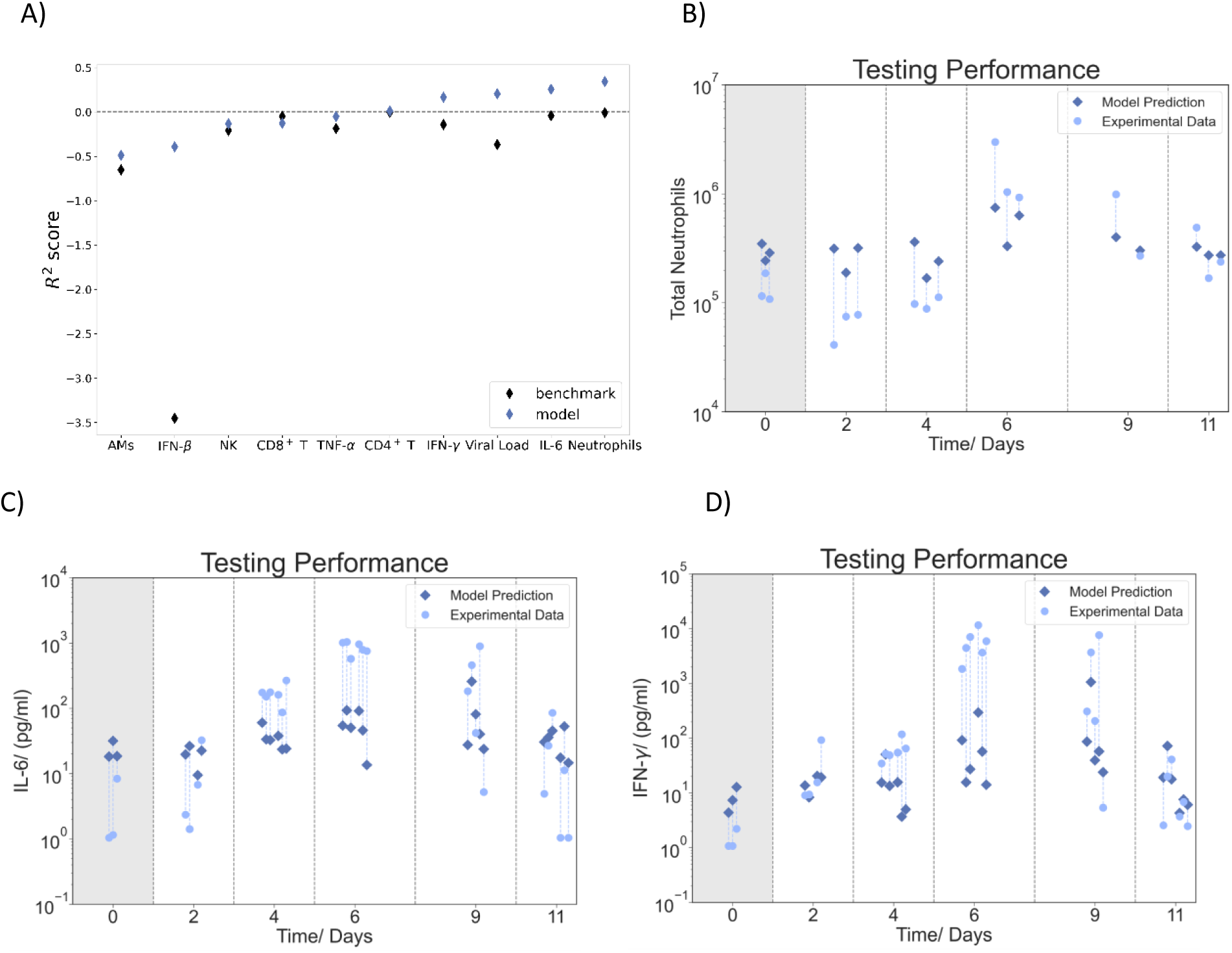
Summary of model predictions for various lung leukocytes and cytokines from hematological data. (A) R^2^ score for different target variables. Blue diamonds show the R^2^ score of the best-performing model. Black diamonds indicate the mean of the target variable obtained from the training data set and serve as a benchmark. Models that perform better than the benchmark and have a positive R^2^ score indicate the model can make successful predictions. Mapping of (B) neutrophils, (C) IL-6, and (D) IFN-γ from blood data. Each blue-light circle represents one mouse, with its matching individual prediction indicated by a connected (blue dashed line) diamond. Gradient boosted regression trees and linear regression with the aid of PCA worked best for these mappings. We observed that the quality of the predictions was dependent on the stage of the infection. Neutrophils were estimated more precisely in the advanced stage of infection, while for IL-6 more accurate estimations were yielded at the peak of the infection. For complete results, please refer to the supplemental material (Supplementary Figures S13-15).

To determine the role of each feature from the hematological data for the prediction of lung outcomes, we performed a feature importance analysis. For this, we calculated the permutation importance by swapping out features and evaluating the performance of our testing data. The results of the feature importance analysis are organized from top to bottom in the level of importance in Figure 5. For example, from all hematological parameters, granulocytes showed the greatest impact on the performance of the machine learning models for predicting the viral load, neutrophils, IFN-γ, and IL-6. Interestingly our analyses revealed a pivotal role of blood platelets for predicting both pathogen burden and lung inflammatory milieu (Figure 5). Granulocytes and erythrocytes ranked second and third place, respectively. The feature importance plots of the additional lung target immune cells and cytokines can be found in the supplemental material (Figure S16).

**Figure 5.**
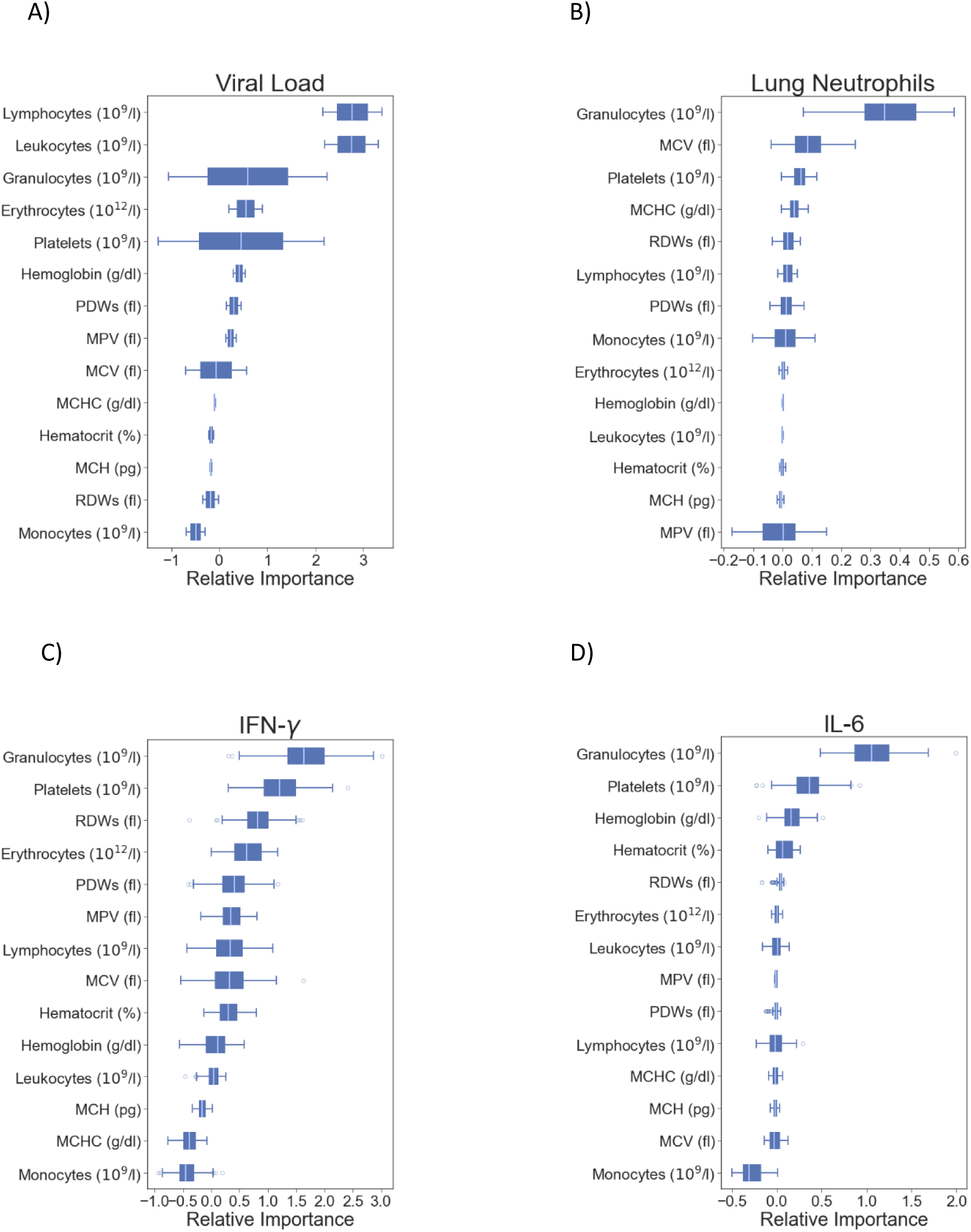
Permutation importance. The permutation importance for the mapping of (A) the viral load, (B) neutrophils, (C) IFN-γ, and (D) IL-6 is indicated. We calculated the permutation importance using the testing data.

## DISCUSSION

Mathematical modeling of host immune responses has largely contributed to improving our understanding of the overall course of influenza infection^28–37^ as well as the personalization of therapies and vaccines^38–40^. Mathematical models consist of systems of ordinary differential equations describing the viral dynamics within the host. However, computational tools for the diagnosis and tracking of respiratory diseases remain a public health challenge.

Here, we progress from the state of the art showing for the first time that minimal-invasively acquired haematological parameters can be used to infer lung viral burden, leukocytes, and cytokines following IAV infection in mice. Nevertheless, despite standardized experimental procedures, our analysis showed a large variance in the computational predictions. These can be attributed to the relatively high variances of our experimental data due to biological or experimental variations. For instance, we found differences in some hematological parameters between the training and the testing experiments, which possibly explain the differences in performance between training and testing prediction of the lung viral load for day 2 post-infection.

The clinical potential of the framework proposed here consists of a new qualitative vision of the disease processes in the lung compartment. We show that the accumulation and decline of multiple cell types involved in the anti-viral immune response in the lung can successfully be predicted with data derived from peripheral blood analyses. The boosted regression tree, with some modifications, provided the best results for many of the lung immune target cells. On the other hand, some target variables proved to be difficult to predict from hematological data. For instance, alveolar macrophage (AM) numbers could not be predicted with any of the tested algorithms and showed the worst score. This is likely the result of the weak correlations of AMs with the different blood cells analyzed. AMs also demonstrate a quite different behavior than the other cells in terms of their abundance over the course of infection as their number peaked rather late during the infection, *i*.*e*. when the virus was mostly cleared^41^. This phenomenon is most likely the result of inflation of the alveolar macrophage pool by self-renewal^42^ and/or monocyte recruitment processes^43^. Regarding the less accurate predictions of cytokines within the airways from blood parameters (Supplementary Figure S13), a possible explanation is that a large portion of these cytokines (especially during the early infection stage) originated from lung-resident leukocytes as well as non-leukocytes. Therefore, their temporal quantity and composition are largely determined by local constituents of the lung’s immune cell response.

Our results show an active reaction chain between peripheral blood parameters and immune cells in the lungs of the mice following IAV infection. Interestingly, we found that peripheral blood platelets play an important role in predicting lung immune cell numbers in IAV infection. In line with our finding of increased numbers of platelets in the blood during the acute phase of IAV infection (see Supplementary Figure S1), platelet accumulation in the pulmonary capillaries is a hallmark of murine IAV H1N1 infection^44^ and contributes to pathogenesis^45^. Importantly, platelet-derived cytokines such as IL-1β can directly increase endothelial permeability and the expression of important vascular adhesion molecules^46,47^. In line with this, airway IL-1β levels were elevated during acute IAV infection (see Figures S8-9). Increased platelet-mediated transendothelial migration of CD4^+^ T cells, CD8^+^ T cells, NK cells, and neutrophils could thus be one conceivable mechanism contributing to the observed strong positive correlation between peripheral blood platelets and the aforementioned lung leukocyte subsets. This is relevant to the anti-viral host response, as the CD4^+^ T cells are central in the activation and maturation of virus-specific CD8^+^ T cells^48^, while neutrophils are required for proper NK cell maturation^49^.

It should be noted, that although the estimation is called “prediction” in the machine learning domain, and blood-derived data can here be used in a practical way to “predict” the viral load or the number of certain lung immune cells or cytokines in IAV infection, we cannot establish a direction of causality in this case. In other words, we cannot state *e*.*g*. that platelets are involved in raising the total amount of CD8^+^ T cells or if CD8^+^ T cells drive the increase in platelets. What we can learn from the correlations between variables is that CD8^+^ T cells, CD4^+^ T cells, NK cells, and neutrophils have their strongest positive correlation with platelets between the blood cells analyzed (see Figure 2). The feature importance analysis confirms that platelets play the most important part in the estimation of lung CD4^+^ T and CD8^+^ T cells, followed by erythrocytes. However, the viral load inside the lungs has also a strong correlation with platelets but an even stronger one with granulocytes. The variable importance analysis suggests that platelets and granulocytes do not strongly contribute to the prediction as can be seen in Figure 5.

The weak predictive results obtained with the linear regression model signifies that these relations have a high order of complexity. While contributing to the viral clearance, the innate immune system can also exacerbate the lung injury^50,51^. In this context, tissue injury can be a cause of platelet activation during influenza infection. The role of platelets in human influenza infection has been stressed in recent years^45,52,53^. Thrombosis, controlled by the innate immune system has been suggested to support immune defense^54^.

Hematological parameters such as neutrophil, lymphocyte, and platelet counts, as well as the neutrophil-to-lymphocyte ratio (NLR) have contributed to diagnosing influenza virus infections^52^. Thus, we also addressed if the use of the granulocyte-lymphocyte ratio (GLR) or the platelet-granulocyte ratio (PGR) improves importance in our model predictions (Table S2). We compared the use of only GLR or PGR with the only use of lymphocytes and granulocytes, as well using additional important IAV infection-associated peripheral blood parameters like erythrocytes and hemoglobin. Performances were evaluated with the MSE and R^2^ scores on the testing data set. We also calculated the corrected Akaike Information Criterion (AIC_c_) during the training to take the complexity of the models into account. Supplementary Table S2 shows that the model performance increased using GLR, while the use of PGR had only a minor effect. Also, adding erythrocytes and hemoglobin did not improve predictions.

In summary, blood platelets, granulocytes, and erythrocytes play an important role in understanding the immune response to influenza infection and can be used in conjunction with other blood components for monitoring the lung viral load and lung immune cells in mice. Importantly, our results indicate that a reduced number of variables does not affect model/prediction accuracy. This can help to further reduce the hematology data needed for successful prediction. While recent efforts show evidence for the diagnosis of COVID-19 from blood compartment^56^, further clinical evidence will be needed to show the potential of how our procedure could be generalized to advance medical care.

## METHODS

### Experimental Design

Mice were intranasally infected with a sublethal dose of the mouse-adapted, strictly pneumotropic H1N1 IAV strain PR/8/34 and the lung viral burden, pulmonary innate and adaptive leukocyte subsets, pulmonary cytokine levels, and peripheral blood cell parameters were assessed for 11 days pi (Figure 1).

Initial generation, training, and optimization of the computational algorithms were conducted using data from two independent *in vivo* infection experiments. Here, mice were randomly assigned to the respective experimental groups and were either intranasally inoculated with a sublethal dose of IAV or saline (control groups). Mice (*n* = 4-6/experimental group) were sacrificed on days 1, 2, 3, 4, 5, 7, 9 and 11 post-infection. Experimental readouts for the first experiment were: hematological parameters and lung tissue viral load. In the second experiment, experimental readouts were: hematological parameters, leukocytes in lung tissue, and airway cytokines.

For subsequent model validation, two additional, independent *in vivo* infection experiments were performed using the above-mentioned readouts. In these experiments, mice (*n* = 3/experimental group) were sacrificed at days 2, 4, 6, 9, and 11 post-infection. Hematological parameters used for model generation and evaluation are listed in Table 1.

### Mice

For all experiments, female C57BL/6JOlaHsd mice (age 10-12 weeks) from Envigo were used. All mice were housed in the animal facility at the Helmholtz Centre for Infection Research under specific-pathogen-free (SPF) conditions and in accordance with national and institutional guidelines.

### Viral preparation and infection

For viral infections, a mouse-adapted influenza A virus strain (A/Puerto Rico/8/34, H1N1) was utilized. The virus was produced in Madin-Darby Canine Kidney (MDCK) cells^57^ and quantified by calculating the tissue culture infectious dose (TCID50) as previously described^58^. Mice were anesthetized by intraperitoneal injection of ketamine/xylazine and were infected with viral inoculum (0.31 TCID50 in 25µL PBS). Control animals received PBS only.

### Quantification of the lung viral load

Lungs were perfused using PBS. RNA was extracted from whole lung tissue homogenates using the RNeasy Plus Mini Kit (Qiagen). The absence of genomic DNA in RNA samples was initially confirmed by PCR using a Taq DNA-polymerase and primers for the housekeeping gene Rps9. Quantitative real-time RT-PCR (qPCR) for detection of viral burden was performed using the SensiFASTTM SYBR® No-ROX One-Step Kit and an influenza nucpleoprotein (NP) plasmid standard. The sequences of the used primers were: 5 ′ CTGGACGAGGGCAAGATGAAGC, 3′TGACGTTGGCGGATGAGCACA (*Rps9*) and 5′ GAGGGGTGAGAATGGACGAAAAAC, 3′CAGGCAGGCAGGCAGGACTT (*Np*).

### Hematology analysis

Blood samples were obtained from the retrobulbar plexus and EDTA was added to prevent coagulation. Samples were analyzed using a VetScan® HM5 machine (Abaxis).

### Cytokine detection

Bronchoalveolar lavage (BAL) was performed with 1mL PBS, samples were spun down (420 x g, 10 min) and BAL fluid (BALF) supernatants were stored at -70°C until further analyses. Cytokine levels in BALF samples were quantified using the LEGENDplex™ Mouse Inflammation Panel (BioLegend) according to the manufacturer’s protocol.

### Isolation of leukocytes from lung tissue

Lungs were perfused using PBS, excised, mechanically homogenized, and enzymatic digestion was performed in Iscove’s modified Dulbecco’s medium (IMDM) supplemented with 0.2mg/mL collagenase D (Roche), 0.01mg/mL DNase I (Sigma-Aldrich), and 5% fetal bovine serum at 37°C for 45min. Digestion was stopped by the addition of EDTA, the cell suspension was filtered through a 100µm cell strainer and spun down (420 x g, 10min). Erythrocyte lysis was performed using ammonium-chloride-potassium (ACK) buffer and leukocytes were isolated by gradient centrifugation using Percoll solution (GE Healthcare). Lung tissue leukocytes were filtered again and antibody staining for flow cytometry was performed.

### Flow cytometry

Lung tissue leukocyte samples were subjected to viability staining and blocking of Fc-receptors using LIVE/DEAD™ Fixable Blue Dead cell stain kit (life technologies) and an anti-CD16/32 antibody (clone 93, BioLegend). Cells were then washed and incubated with a staining mix containing antibodies against the following murine antigens: Siglec-F (PE, clone E50-2440, BD), Ly6G (AlexaFluor700, clone 1A8, BD), CD11c (APC, clone N 418, BioLegend), CD11b (BV421, clone M1/70, BD), CD4 (APC-Fire750, clone GK1.5, BioLegend), CD8a (CyChrome, clone 53-6.7, BD), CD3ε (Biotin, clone 145-2C11, BioLegend), NK1.1 (FITC, clone PK136, BioLegend). Secondary staining was performed using streptavidin-BV605 and streptavidin-BV650, respectively (BioLegend). All reagents and antibodies had been titrated before the experiments for optimal staining results. Flow cytometry data were acquired using LSRII and LSR Fortessa instruments (BD). Data were analyzed using FlowJo software (BD).

### Data Processing

The data obtained from the hematological analysis constitutes of 20 different parameters. Some parameters are given in absolute values and percentages. We considered from these parameters only the absolute values, resulting in 14 different parameters (see Table 1). For some mice, it was not possible to extract hematological data and/or according to target infection marker. These mice were removed for the mapping, although their data was used in the correlation analysis. Furthermore, if some values were lower than the measurable threshold, we used the threshold value. Computational algorithms were implemented in python using the *Keras* and *sklearn* libraries. Our study design yielded separate training and testing data sets. The testing data was obtained approximately one year after the training data. All laboratory conditions were kept as similar as possible.

Using the data directly for training the algorithms led to poor results and therefore we used data pre-processing techniques. To conserve the nature of hematological parameter distributions, we used the min-max scaling from the sklearn.preprocessing. MinMaxScaler class:

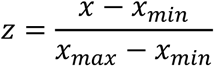

For the target infection markers like viral load, any logarithmic function worked well. For simplicity, we used log_10_.

### Machine Learning Models

Different machine learning models were tested for the mapping including feedforward neural networks (FNN), gradient boosted regression trees (GBRT), linear regression (LR), support vector machines (SVM), and random forest regression (RFR). The hyperparameters of the models were estimated via grid-search and adjusted via trial and error. FNN and GBRT showed to superior in most cases and RFR was outperformed in every instance with one of the other algorithms.

In many cases using PCA before the mapping yielded improved performance. It was found that a dimensionality reduction to six input blood variables was often best. We used the class *sklearn*.*decomposition*.*PCA* for implementation. For the feedforward neural network, the *keras* library was used with a TensorFlow backend. We found that one hidden layer was sufficient most of the time and additional layers were not needed. The number of weights varied from 10 to 50. For regularization, the addition of dropout layers with a rate of 0.2 was helpful to prevent overfitting. As an activation function, we used a rectified linear unit (ReLU). We had one output that uses a linear activation function. This was necessary to map the whole range of possible outcome values. We used the Adam optimizer and minimized the mean squared error to find the optimal fit. The weights were initialized according to a He-uniform distribution.^59^ Following common practice in literature we used for the training of the neural network a validation set of 10% of the whole training data.

The GBRT, LR, SVM, and RFR algorithms are taken from the python library *sklearn*. The hyperparameters of GBRT and RFR models were searched over a grid from 10 to 2000 estimators, a learning rate from 0.001 to 0.09, and a max depth of 2 to 14. The least-square regression was used for optimization. The kernels used for SVM were *‘linear’, ‘poly’, ‘rbf ‘, ‘sigmoid’* and *‘precomputed’*.

To determine which variables were the most important in our model predictions, we calculated the permutation importance using the sklearn.inspection.permutation_importance implementation. For this, we took our best model, respectively, and trained it on the training data set. After the trained model was evaluated on the hold-out testing data set with the mean squared error as metric, a feature column was permuted and the metric was evaluated again. This procedure was repeated 100 times and the permutation importance was given by the difference between the baseline metric and the metric from permutated feature columns.

## DATA AND CODE AVAILABILITY

The datasets generated and analyzed during the current study are available: https://github.com/Jhutty/Tracking_IAV_from_Blood

## ACKNOWLEDGEMENTS

We thank Tatjana Hirsch, Hanna Shkarlet and Karin Lammert for expert technical assistance in infection experiments. This work was supported by the Deutsche Forschungsgemeinschaft with the project HE7707/5-1 and BR2221/6-1; the Universidad Nacional Autonoma de Mexico (UNAM) – PAPIIT with the number IA102521; and the Alfons und Gertrud Kassel-Stiftung.

## ETHICS DECLARATIONS

All the experiments were approved and conducted in accordance with the guidelines set by the local animal welfare and ethics committee (Niedersächsisches Landesamt für Verbraucherschutz und Lebensmittelsicherheit).

## REFERENCES

1. World Health Organization. Influenza (Seasonal) fact sheet.

2. WHO Recommended Surveillance Standards. Second edition.

3. Kilbourne, E. D. Influenza Pandemics of the 20th Century. Emerging Infectious Diseases 12, 9 (2006).

4. World Health Organization. Writing Committee of the WHO Consultation on Clinical aspects of pandemic 2009 influenza A (H1N1) virus infection. New England Journal of Medicine 1708–1719 (2010).

5. Sharma-Chawla, N. et al. In vivo neutralization of pro-inflammatory cytokines during secondary streptococcus pneumoniae infection post influenza a virus infection. Frontiers in Immunology 10, (2019).

6. Allwinn, R. et al. Laboratory diagnosis of influenza – virology or serology? Medical Microbiology and Immunology 2002 191:3 191, 157–160 (2002).

7. L, D. et al. Cellular changes in blood indicate severe respiratory disease during influenza infections in mice. PloS one 9, (2014).

8. IE, G. et al. Untuned antiviral immunity in COVID-19 revealed by temporal type I/III interferon patterns and flu comparison. Nature immunology 22, 32–40 (2021).

9. Y, Z. et al. Pathway mapping of leukocyte transcriptome in influenza patients reveals distinct pathogenic mechanisms associated with progression to severe infection. BMC medical genomics 13, (2020).

10. BM, C. et al. Inflammatory Monocytes Drive Influenza A Virus-Mediated Lung Injury in Juvenile Mice. Journal of immunology (Baltimore, Md. : 1950) 200, 2391–2404 (2018).

11. N, S.-C. et al. Influenza A Virus Infection Predisposes Hosts to Secondary Infection with Different Streptococcus pneumoniae Serotypes with Similar Outcome but Serotype-Specific Manifestation. Infection and immunity 84, 3445–3457 (2016).

12. Duvigneau, S. et al. Hierarchical effects of pro-inflammatory cytokines on the post-influenza susceptibility to pneumococcal coinfection. Scientific Reports 6, 1–11 (2016).

13. Ian Goodfellow, Yoshua Bengio & Aaron Courville. Deep learning. MIT press (2016) doi:10.1007/S10710-017-9314-Z.

14. Deng Li & Yu Dong. Deep Learning. Foundations and Trends in Signal Processing 7, 197–387 (2014).

15. Liu, F., Li, H., Ren, C., Bo, X. & Shu, W. PEDLA: predicting enhancers with a deep learning-based algorithmic framework. Scientific Reports 2016 6:1 6, 1–14 (2016).

16. Wang, S., Peng, J., Ma, J. & Xu, J. Protein Secondary Structure Prediction Using Deep Convolutional Neural Fields. Scientific Reports 2016 6:1 6, 1–11 (2016).

17. Quang, D., Chen, Y. & Xie, X. DANN: a deep learning approach for annotating the pathogenicity of genetic variants. Bioinformatics 31, 761–763 (2015).

18. Esteva, A. et al. Dermatologist-level classification of skin cancer with deep neural networks. Nature 2017 542:7639 542, 115–118 (2017).

19. Huang, Q., Zhang, F. & Li, X. Machine Learning in Ultrasound Computer-Aided Diagnostic Systems: A Survey. BioMed Research International 2018, (2018).

20. Kabak, M., Çil, B. & Hocanlı, I. Relationship between leukocyte, neutrophil, lymphocyte, platelet counts, and neutrophil to lymphocyte ratio and polymerase chain reaction positivity. International Immunopharmacology 93, 107390 (2021).

21. Han, Q. et al. Role of hematological parameters in the diagnosis of influenza virus infection in patients with respiratory tract infection symptoms. Journal of Clinical Laboratory Analysis 34, e23191 (2020).

22. Hernandez-Vargas, E. A. et al. Effects of Aging on Influenza Virus Infection Dynamics. Journal of Virology 88, 4123–4131 (2014).

23. Hernandez-Vargas, E. A. & Velasco-Hernandez, J. X. In-host Mathematical Modelling of COVID-19 in Humans. Annual Reviews in Control 50, 448–456 (2020).

24. Pawelek, K. A. et al. Modeling Within-Host Dynamics of Influenza Virus Infection Including Immune Responses. PLOS Computational Biology 8, e1002588 (2012).

25. Dobrovolny, H. M., Gieschke, R., Davies, B. E., Jumbe, N. L. & Beauchemin, C. A. A. Neuraminidase inhibitors for treatment of human and avian strain influenza: A comparative modeling study. Journal of Theoretical Biology 269, 234–244 (2011).

26. Baccam, P., Beauchemin, C., Macken, C. A., Hayden, F. G. & Perelson, A. S. Kinetics of Influenza A Virus Infection in Humans. Journal of Virology 80, 7590–7599 (2006).

27. Smith, A. M. et al. Kinetics of Coinfection with Influenza A Virus and Streptococcus pneumoniae. PLOS Pathogens 9, e1003238 (2013).

28. Miao, H., Xia, X., Perelson, A. S. & Wu, H. On Identifiability of Nonlinear ODE Models and Applications in Viral Dynamics. SIAM Review 53, 3–39 (2011).

29. Canini, L. & Perelson, A. S. Viral kinetic modeling: state of the art. Journal of Pharmacokinetics and Pharmacodynamics 41, 431–443 (2014).

30. Canini, L. & Carrat, F. Population modeling of influenza A/H1N1 virus kinetics and symptom dynamics. Journal of Virology 85, 2764–2770 (2011).

31. Handel, A. & Antia, R. A simple mathematical model helps to explain the immunodominance of CD8 T cells in influenza A virus infections. Journal of virology 82, 7768–72 (2008).

32. Beauchemin, C. & Handel, A. A review of mathematical models of influenza A infections within a host or cell culture: lessons learned and challenges ahead. BMC public health 11, S7 (2011).

33. Hancioglu, B., Swigon, D. & Clermont, G. A dynamical model of human immune response to influenza A virus infection. Journal of Theoretical Biology 246, 70–86 (2007).

34. Smith, A. M. Host-pathogen kinetics during influenza infection and coinfection: insights from predictive modeling. Immunological Reviews vol. 285 97–112 (2018).

35. Smith, A. M. & Perelson, A. S. Influenza A virus infection kinetics: quantitative data and models. Wiley interdisciplinary reviews. Systems biology and medicine 3, 429–445 (2011).

36. Baccam, P., Beauchemin, C., Macken, C. a, Hayden, F. G. & Perelson, A. S. Kinetics of influenza A virus infection in humans. Journal of virology 80, 7590–9 (2006).

37. Harper, S. A. et al. Seasonal Influenza in Adults and Children—Diagnosis, Treatment, Chemoprophylaxis, and Institutional Outbreak Management: Clinical Practice Guidelines of the Infectious Diseases Society of America. Clinical Infectious Diseases 48, 1003–1032 (2009).

38. Hernandez-Mejia, G. & Hernandez-Vargas, E. A. Uncovering antibody cross-reaction dynamics in influenza A infections. bioRxiv 2020.01.06.896274 (2020) doi:10.1101/2020.01.06.896274.

39. Hernandez-Mejia, G., Alanis, A. Y. & Hernandez-Vargas, E. A. Inverse Optimal Impulsive Control Based Treatment of Influenza Infection. in IFAC World Congress 2017 vol. 50 12696–12701 (2017).

40. Parra-Rojas, C., Messling, V. & Hernandez-Vargas, E. A. Adjuvanted influenza vaccine dynamics. Scientific Reports 9, (2019).

41. Toapanta, F. R. & Ross, T. M. Impaired immune responses in the lungs of aged mice following influenza infection. Respiratory Research 10, 1–19 (2009).

42. Yao, Y. et al. Induction of Autonomous Memory Alveolar Macrophages Requires T Cell Help and Is Critical to Trained Immunity. Cell 175, 1634-1650.e17 (2018).

43. Aegerter, H. et al. Influenza-induced monocyte-derived alveolar macrophages confer prolonged antibacterial protection. Nature immunology 21, 145–157 (2020).

44. Rommel, M. G. E., Milde, C., Eberle, R., Schulze, H. & Modlich, U. Endothelial–platelet interactions in influenza-induced pneumonia: A potential therapeutic target. Anatomia, Histologia, Embryologia 49, 606–619 (2020).

45. Lê, V. B. et al. Platelet Activation and Aggregation Promote Lung Inflammation and Influenza Virus Pathogenesis. https://doi.org/10.1164/rccm.201406-1031OC 191, 804–819 (2015).

46. Rossaint, J., Margraf, A. & Zarbock, A. Role of Platelets in Leukocyte Recruitment and Resolution of Inflammation. Frontiers in Immunology 0, 2712 (2018).

47. Hawrylowicz, C. M., Howells, G. L. & Feldmann, M. Platelet-derived interleukin 1 induces human endothelial adhesion molecule expression and cytokine production. Journal of Experimental Medicine 174, 785–790 (1991).

48. Zens, K. D. & Farber, D. L. Memory CD4 T Cells in Influenza. Current Topics in Microbiology and Immunology 386, 399–421 (2014).

49. Jaeger, B. N. et al. Neutrophil depletion impairs natural killer cell maturation, function, and homeostasis. Journal of Experimental Medicine 209, 565–580 (2012).

50. Herold, S., Becker, C., Ridge, K. M. & Budinger, G. R. S. Influenza virus-induced lung injury: pathogenesis and implications for treatment. European Respiratory Journal 45, 1463–1478 (2015).

51. Kuiken, T., Riteau, B., Fouchier, R. A. M. & Rimmelzwaan, G. F. Pathogenesis of influenza virus infections: the good, the bad and the ugly. Current Opinion in Virology 2, 276–286 (2012).

52. Koupenova, M. et al. The role of platelets in mediating a response to human influenza infection. Nature Communications 2019 10:1 10, 1–18 (2019).

53. Assinger, A. Platelets and Infection – An Emerging Role of Platelets in Viral Infection. Frontiers in Immunology 0, 649 (2014).

54. Engelmann, B. & Massberg, S. Thrombosis as an intravascular effector of innate immunity. Nature Reviews Immunology 2012 13:1 13, 34–45 (2012).

55. Han, Q. et al. Role of hematological parameters in the diagnosis of influenza virus infection in patients with respiratory tract infection symptoms. Journal of Clinical Laboratory Analysis 34, e23191 (2020).

56. Kukar, M. et al. COVID-19 diagnosis by routine blood tests using machine learning. Scientific Reports 2021 11:1 11, 1–9 (2021).

57. Stegemann, S. et al. Increased Susceptibility for Superinfection with Streptococcus pneumoniae during Influenza Virus Infection Is Not Caused by TLR7-Mediated Lymphopenia. (2009) doi:10.1371/journal.pone.0004840.

58. Reed, L. J. & Muench, H. A SIMPLE METHOD OF ESTIMATING FIFTY PER CENT ENDPOINTS. American Journal of Epidemiology 27, 493–497 (1938).

59. He, K., Zhang, X., Ren, S. & Sun, J. Delving Deep into Rectifiers: Surpassing Human-Level Performance on ImageNet Classification.

