## Supplemental Information for "Tracking influenza a virus infection in the lung from hematological data with machine learning"

**Contributions**

### These authors contributed equally: Suneet Singh Jhutty, Julia D. Boehme

[*] These authors jointly supervised this work: Dunja Bruder, Esteban A. Hernandez-Vargas

### 1 SUPPLEMENTAL EXPERIMENTAL DATA INFORMATION

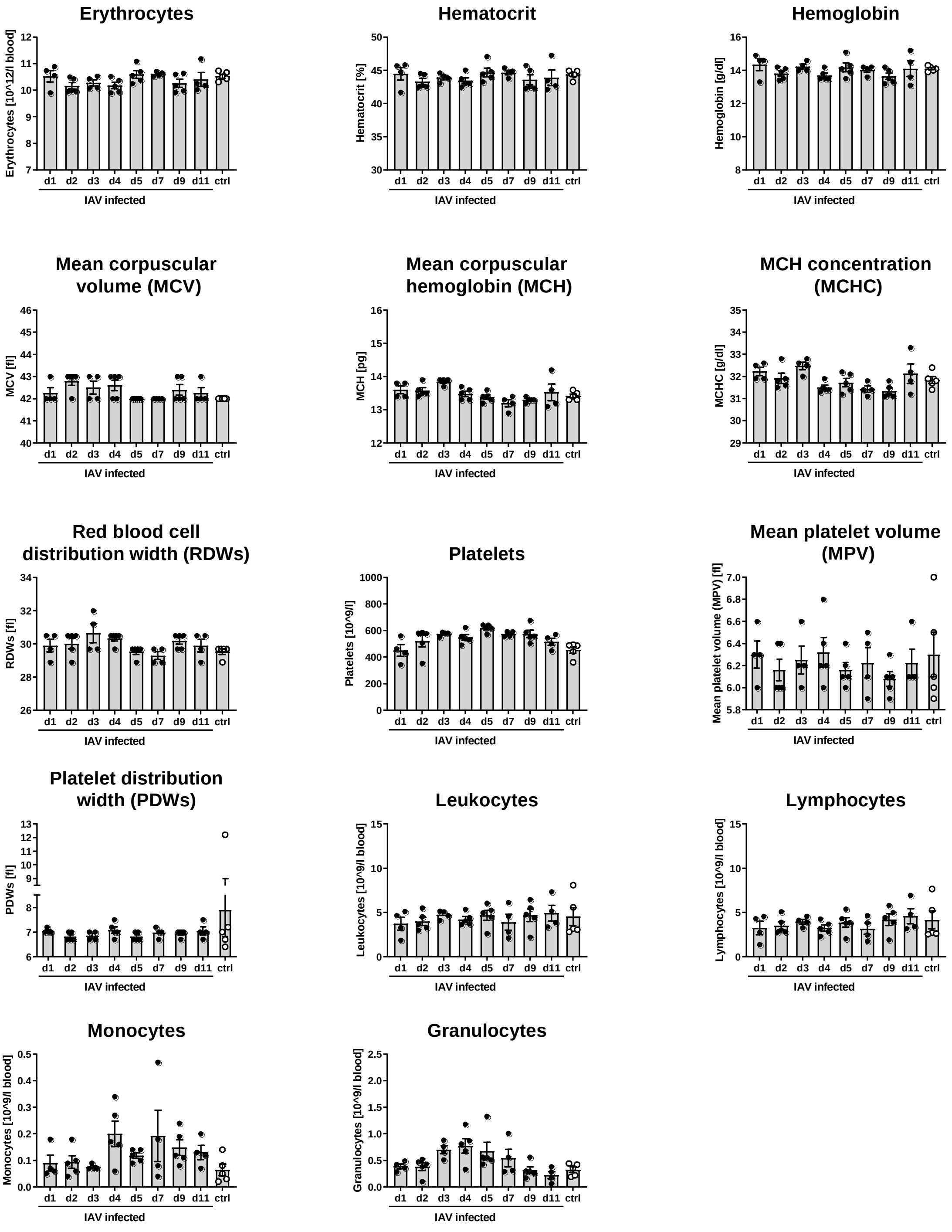


**Figure S1.** Hematological parameters during IAV infection (first experiment, model setup and training). Wild-type C57BL/6JOlaHsd mice were intranasally inoculated with Influenza A virus (IAV) strain A/PR/8/34 (H1N1) or treated with PBS (ctrl) on day 0. At indicated time points, blood was analyzed on a VetScan® HM5 machine. Data for individual mice and mean±SEM are graphed.

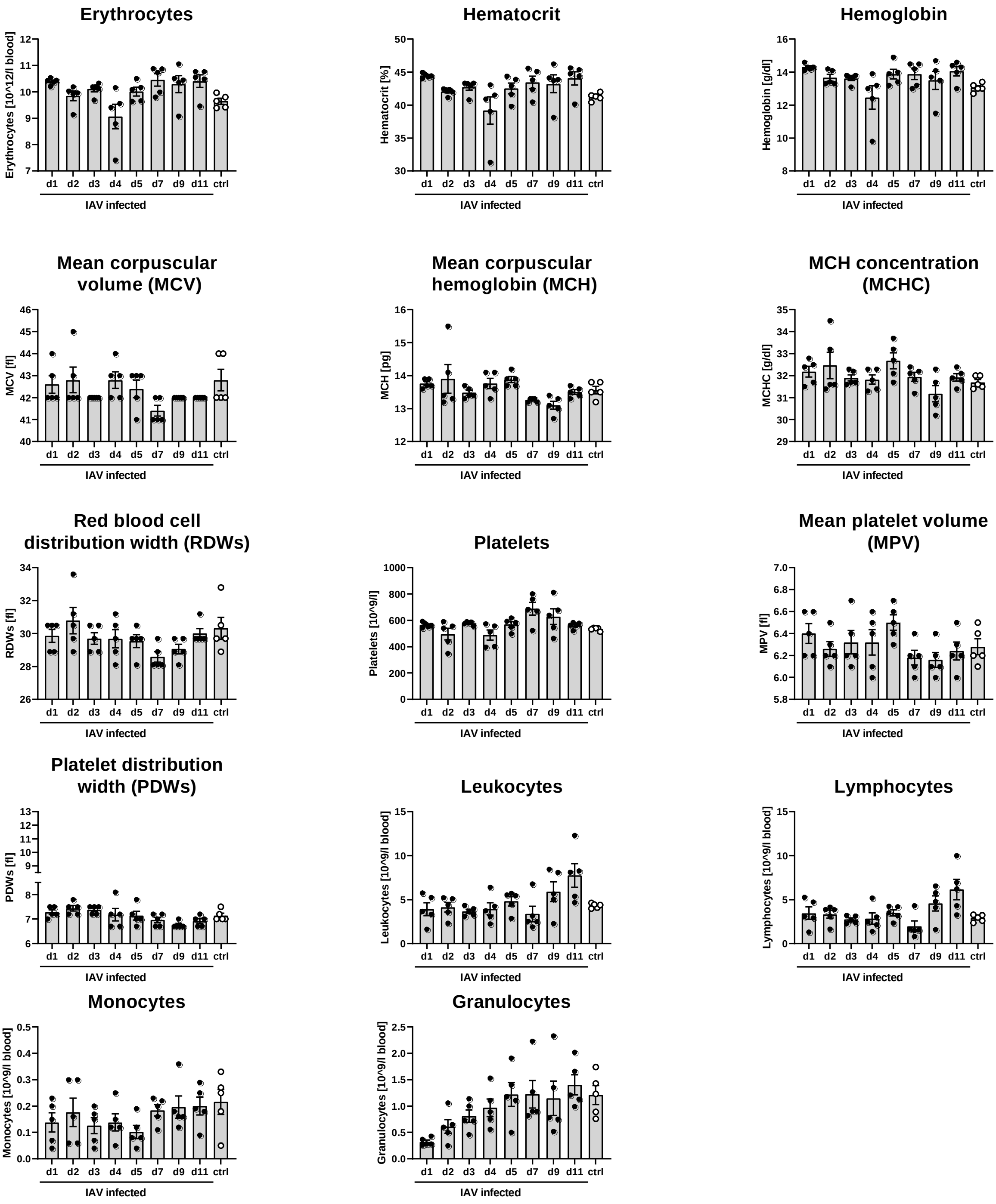


**Figure S2.** Hematological parameters during IAV infection (second experiment, model setup and training). Wild-type C57BL/6JOlaHsd mice were intranasally inoculated with Influenza A virus (IAV) strain A/PR/8/34 (H1N1) or treated with PBS (ctrl) on day 0. At indicated time points, blood was analyzed on a VetScan® HM5 machine. Data for individual mice and mean±SEM are graphed.

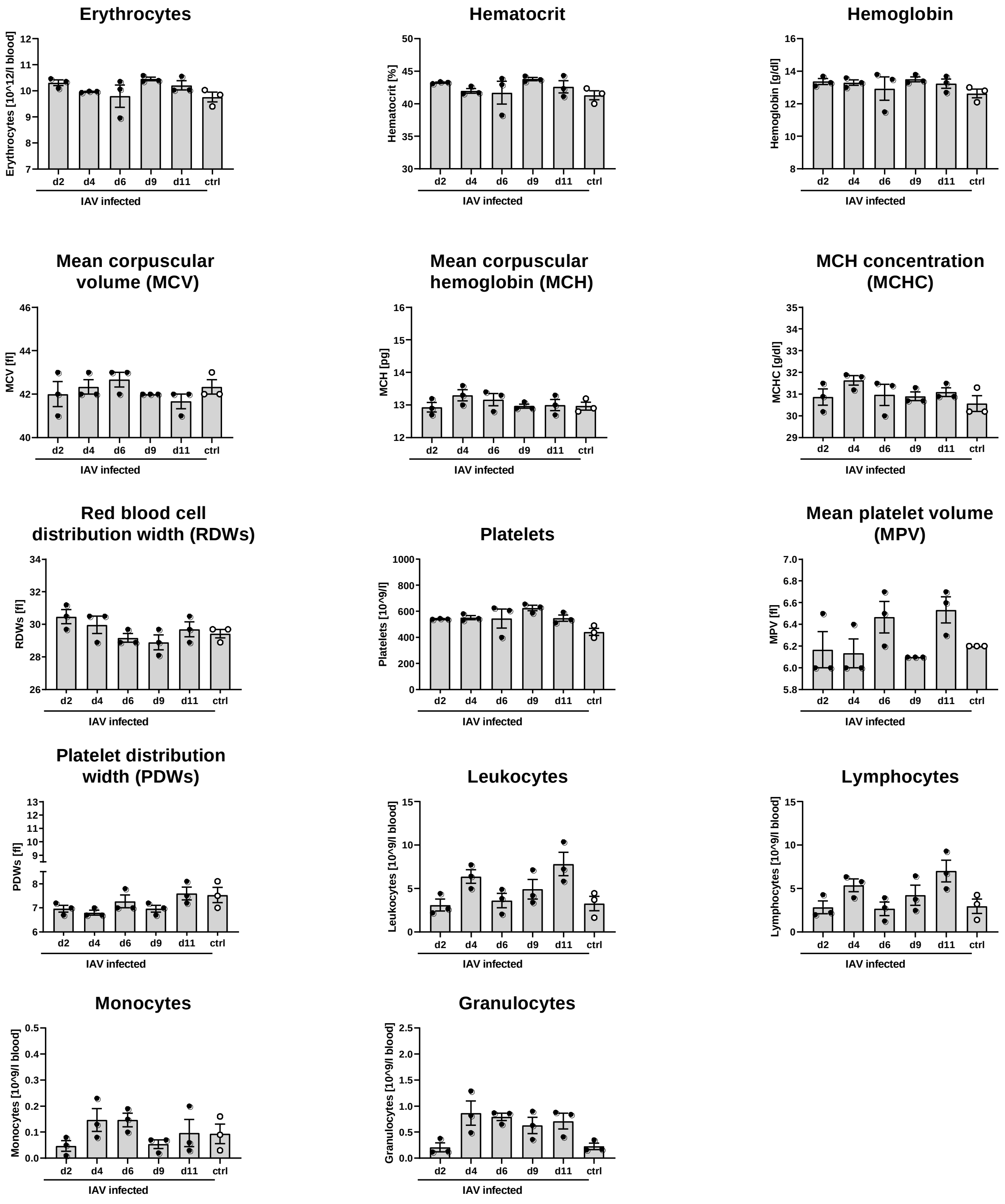


**Figure S3.** Hematological parameters during IAV infection (third experiment, model validation). Wild-type C57BL/6JOlaHsd mice were intranasally inoculated with Influenza A virus (IAV) strain A/PR/8/34 (H1N1) or treated with PBS (ctrl) on day 0. At indicated time points, blood was analyzed on a VetScan® HM5 machine. Data for individual mice and mean±SEM are graphed.

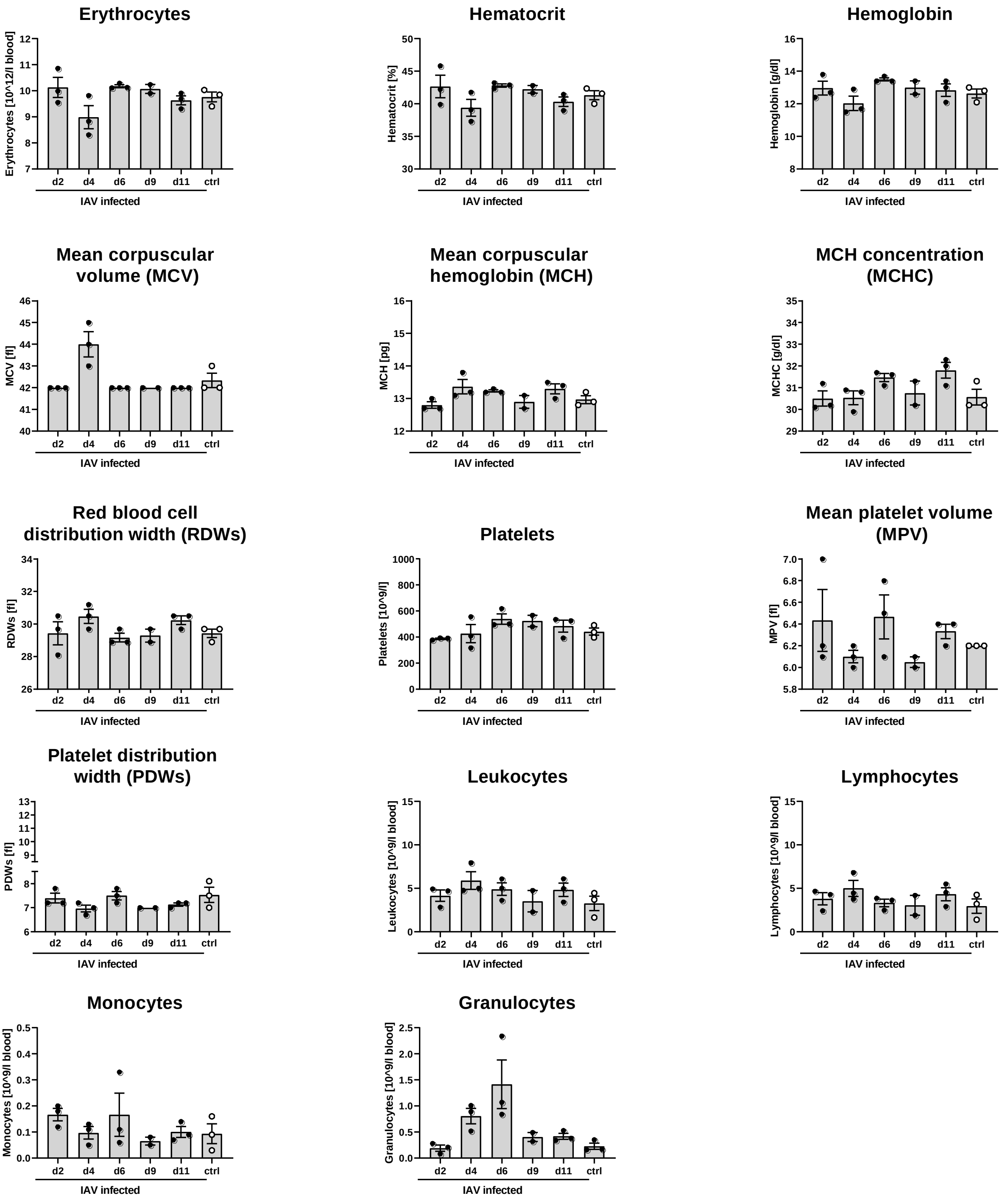


**Figure S4.** Hematological parameters during IAV infection (fourth experiment, model validation). Wild-type C57BL/6JOlaHsd mice were intranasally inoculated with Influenza A virus (IAV) strain A/PR/8/34 (H1N1) or treated with PBS (ctrl) on day 0. At indicated time points, blood was analyzed on a VetScan® HM5 machine. Data for individual mice and mean±SEM are graphed.

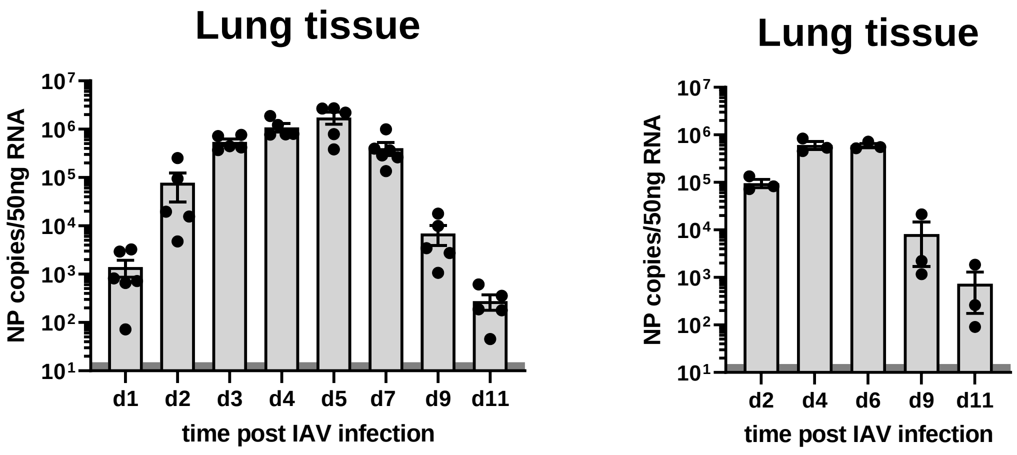


**Figure S5.** Lung viral load during IAV infection. Wild-type C57BL/6JOlaHsd mice were intranasally inoculated with Influenza A virus (IAV) strain A/PR/8/34 (H1N1) on day 0. At indicated time points, RNA from lung tissue homogenates was extracted and viral burden was assessed by quantifying viral nucleoprotein (NP) copy numbers using quantitative real-time RT-PCR. Data for individual mice and mean±SEM are graphed. Dark grey shades indicate the detection limit. Left panel: Data from first experiment (model setup and training). Right panel: Data from third experiment (model testing and evaluation).

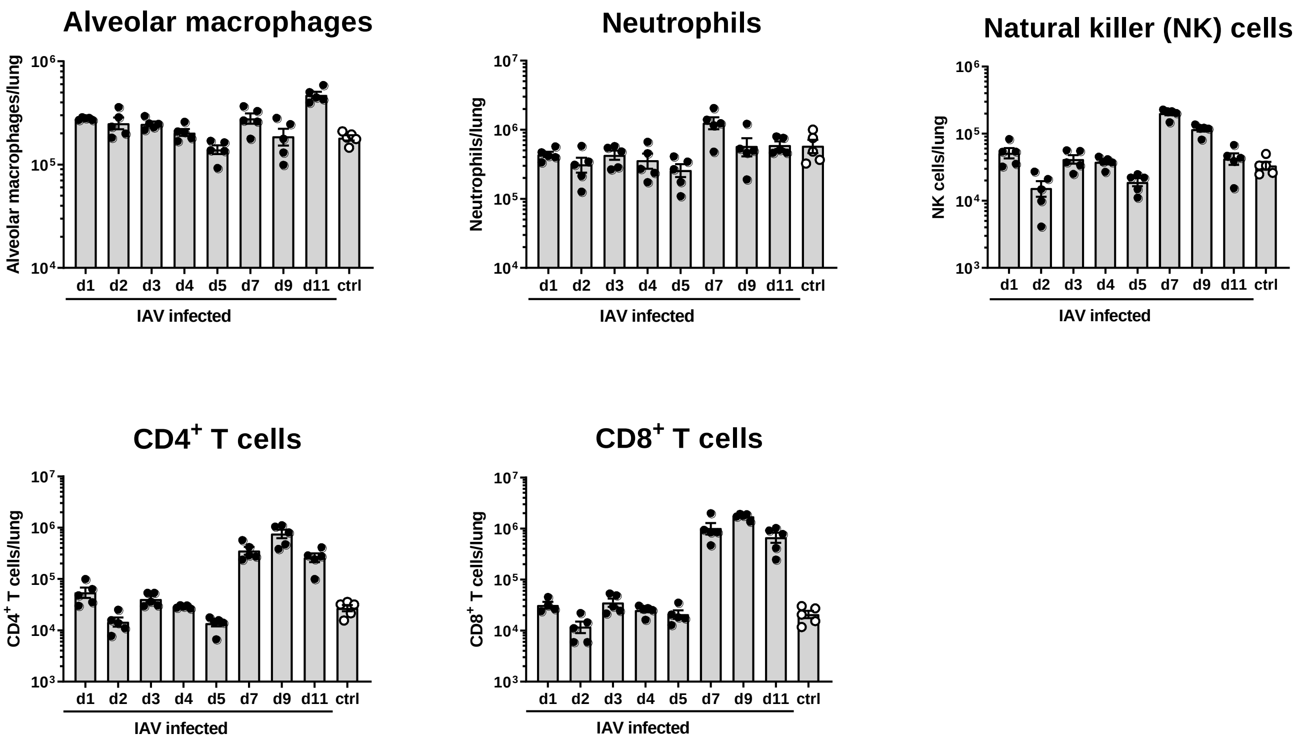


**Figure S6.** Lung leukocytes during IAV infection (second experiment, model setup and training): Wild-type C57BL/6JOlaHsd mice were intranasally inoculated with Influenza A virus (IAV) strain A/PR/8/34 (H1N1) or treated with PBS (ctrl) on day 0. At indicated time points, lung tissue leukocytes were quantified using flow cytometry. Data for individual mice and mean±SEM are graphed.

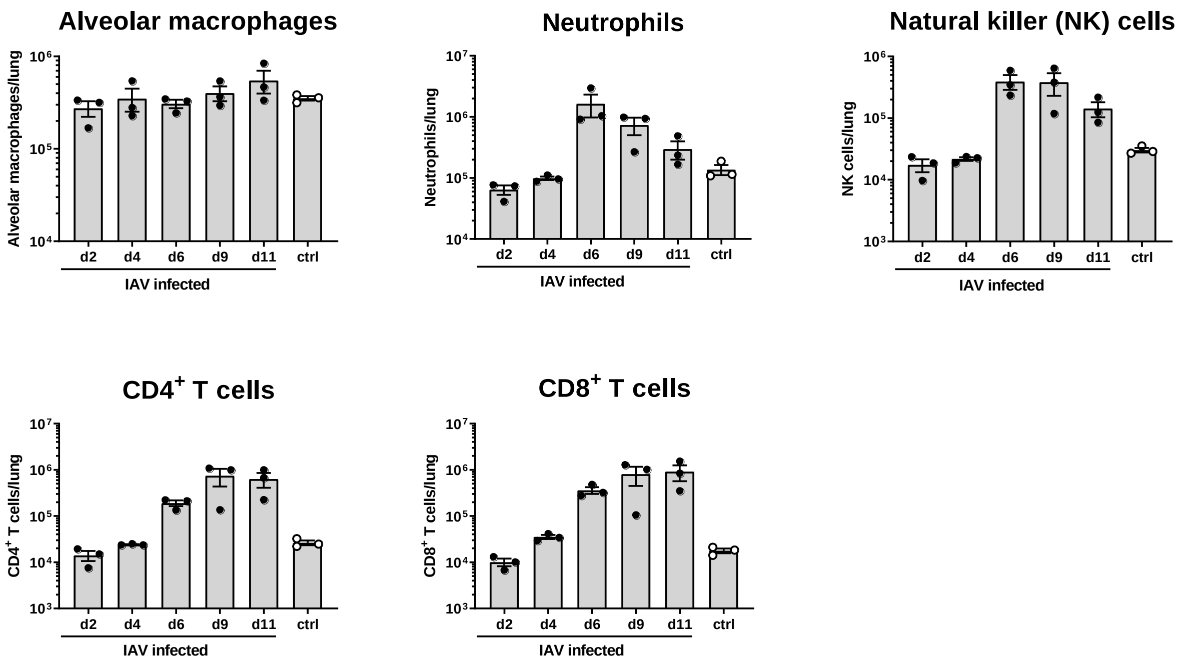


**Figure S7.** Lung leukocytes during IAV infection (fourth experiment, model validation): Wild-type C57BL/6JOlaHsd mice were intranasally inoculated with Influenza A virus (IAV) strain A/PR/8/34 (H1N1) or treated with PBS (ctrl) on day 0. At indicated time points, lung tissue leukocytes were quantified using flow cytometry. Data for individual mice and mean±SEM are graphed.

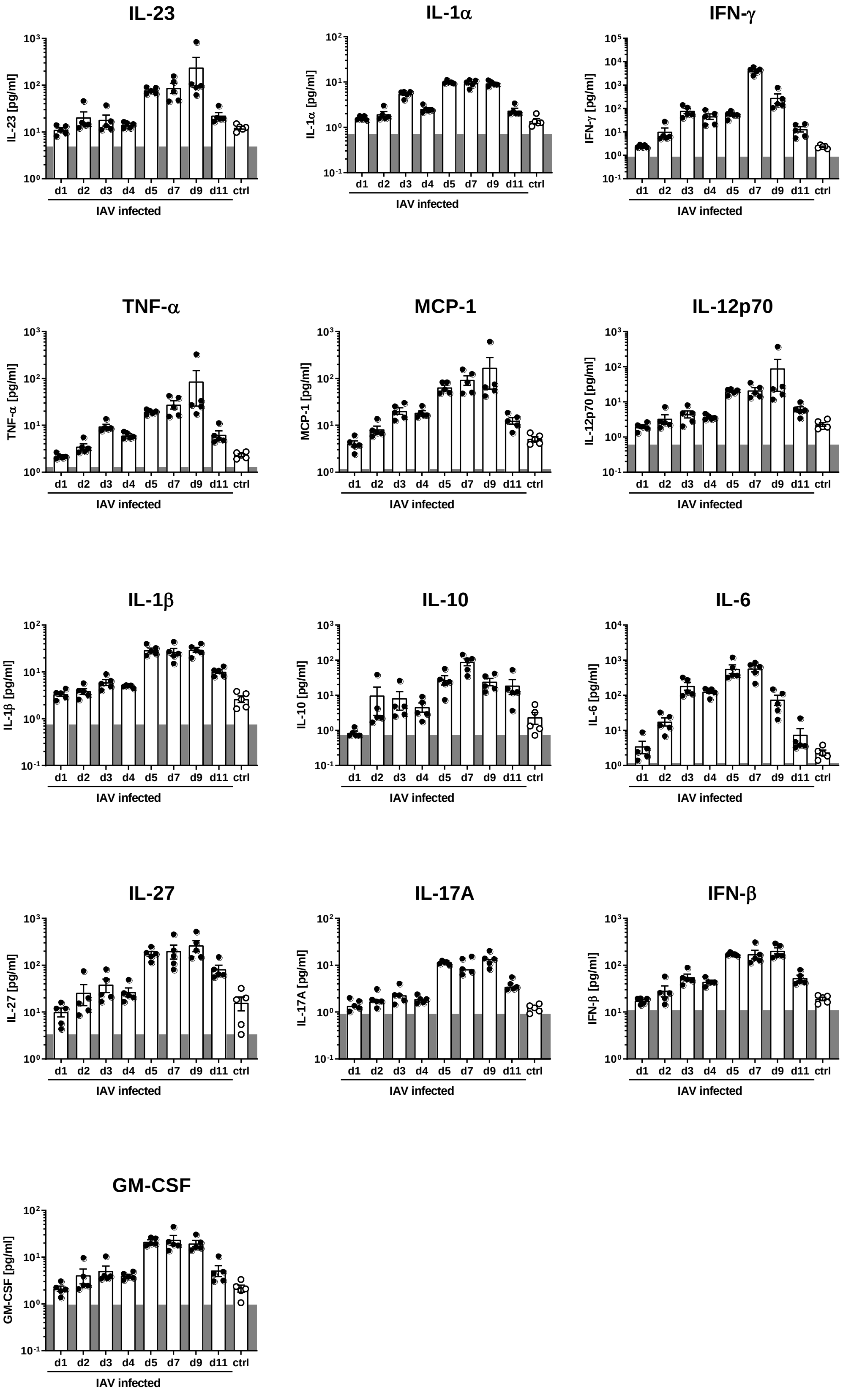


**Figure S8.** Lung cytokines and chemokines during IAV infection (second experiment, model setup and training). Wild-type C57BL/6JOlaHsd mice were intranasally inoculated with Influenza A virus (IAV) strain A/PR/8/34 (H1N1) or treated with PBS (ctrl) on day 0. At indicated time points, concentrations of cytokines and chemokines in the airways were determined by multiplex analyses. Data for individual mice and mean±SEM are graphed. Dark grey shade indicates the detection limit.

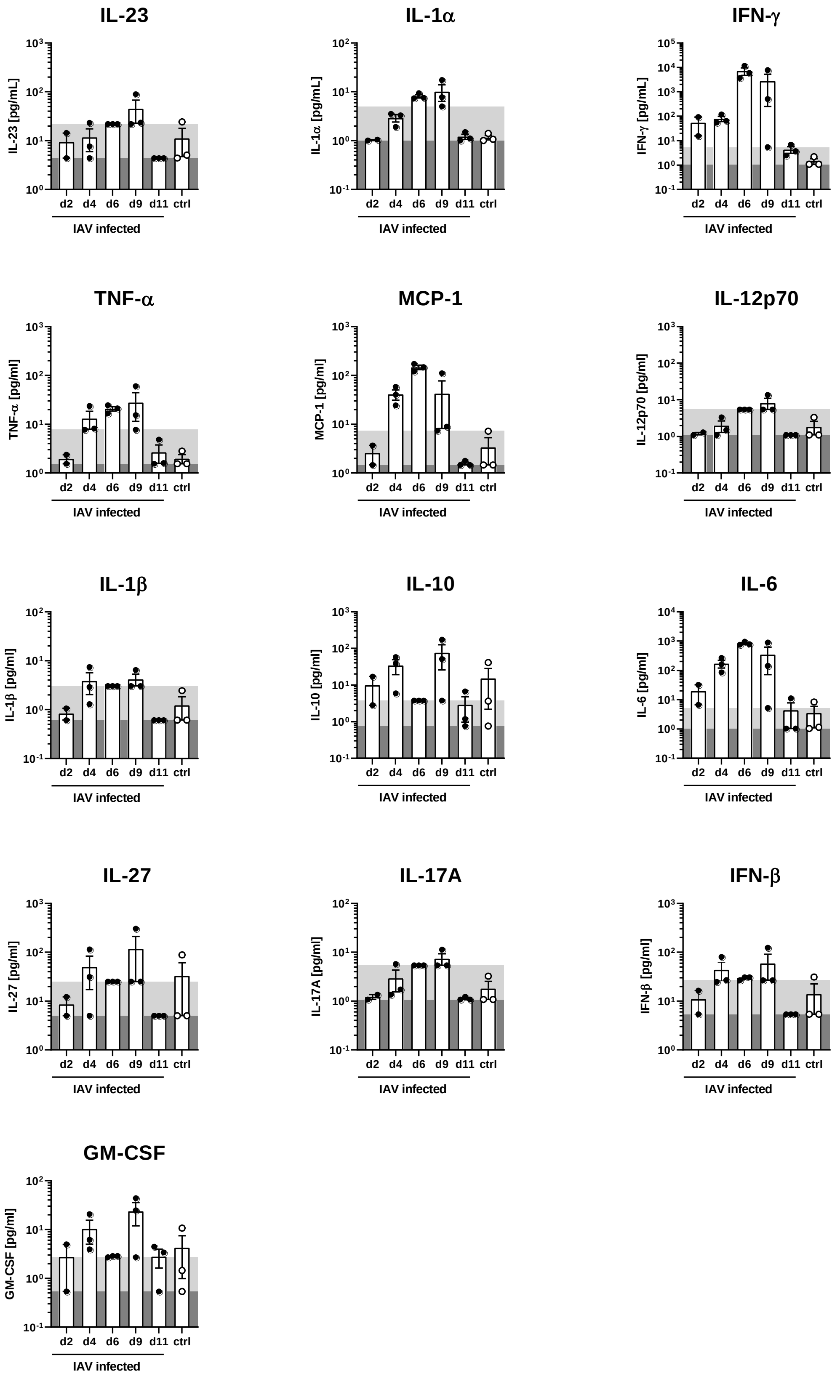


**Figure S9.** Lung cytokines and chemokines during IAV infection (fourth experiment, model validation). Wild-type C57BL/6JOlaHsd mice were intranasally inoculated with Influenza A virus (IAV) strain A/PR/8/34 (H1N1) or treated with PBS (ctrl) on day 0. At indicated time points, concentrations of cytokines and chemokines in the airways were determined by multiplex analyses. Data for individual mice and mean±SEM are graphed. Dark grey shade indicates the detection limit for samples from ctrl, d2, d4 and d11. Light grey shade indicates detection limit for samples from d6 and d9.

#### 2 SUPPLEMENTAL CORRELATION ANALYSIS

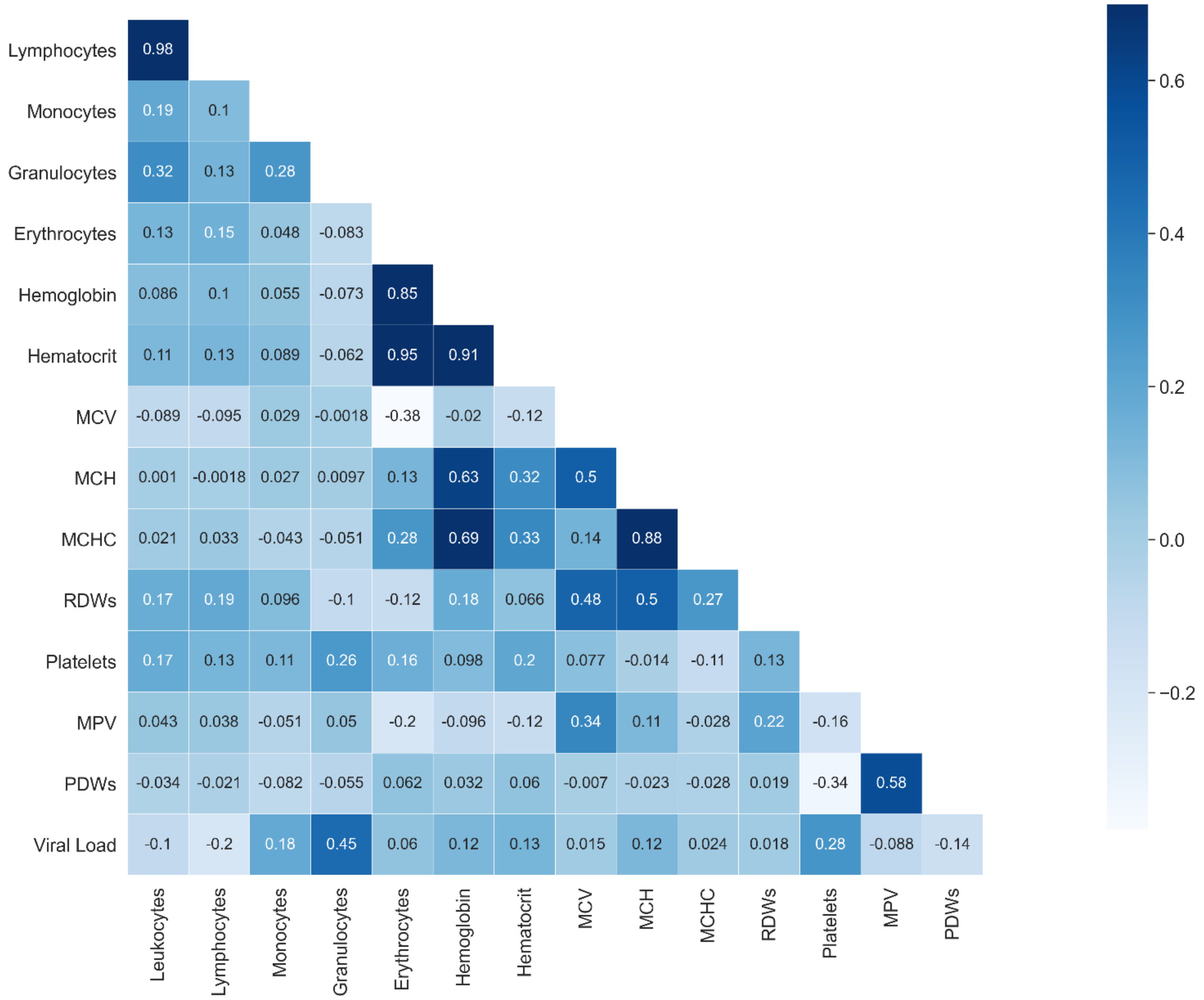


**Figure S10.** Hematological parameters during IAV infection. Correlations of blood cells and viral load over the course of influenza infection. The matrix depicts Pearson correlation coefficients of several different blood cells and the viral load. The data (derived from the first experiment) were used as training data for the machine learning models.

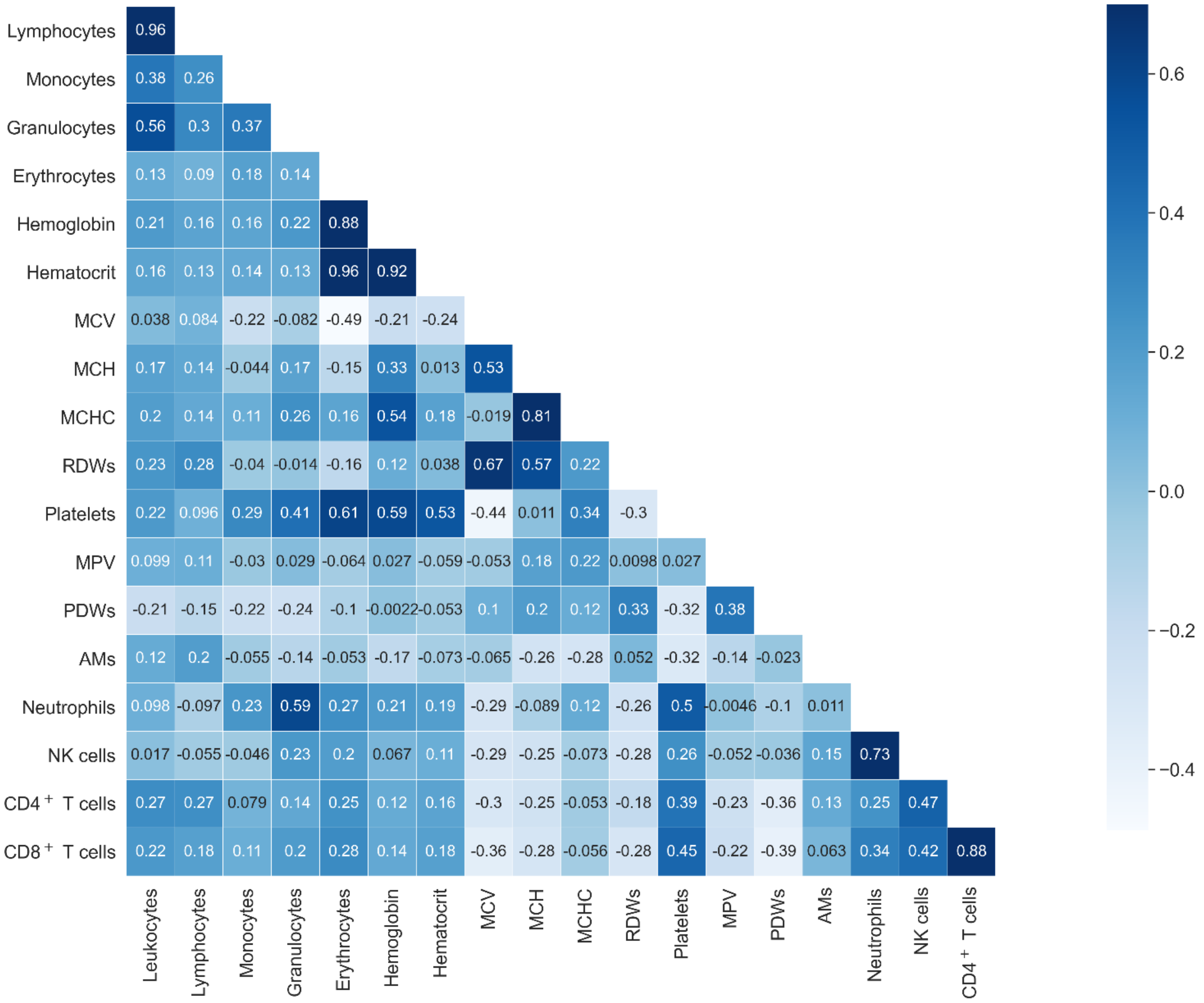


**Figure S11.** Correlations of blood cells and lung leukocytes (alveolar macrophages (AMs), Neutrophils, NK cells, CD4$^{+}$ and CD8$^{+}$ T cells) over the course of influenza infection. The matrix depicts Pearson correlation coefficients of all measured blood cells and lung leukocytes. Data (derived from the second experiment) were used as training data for the machine learning models.

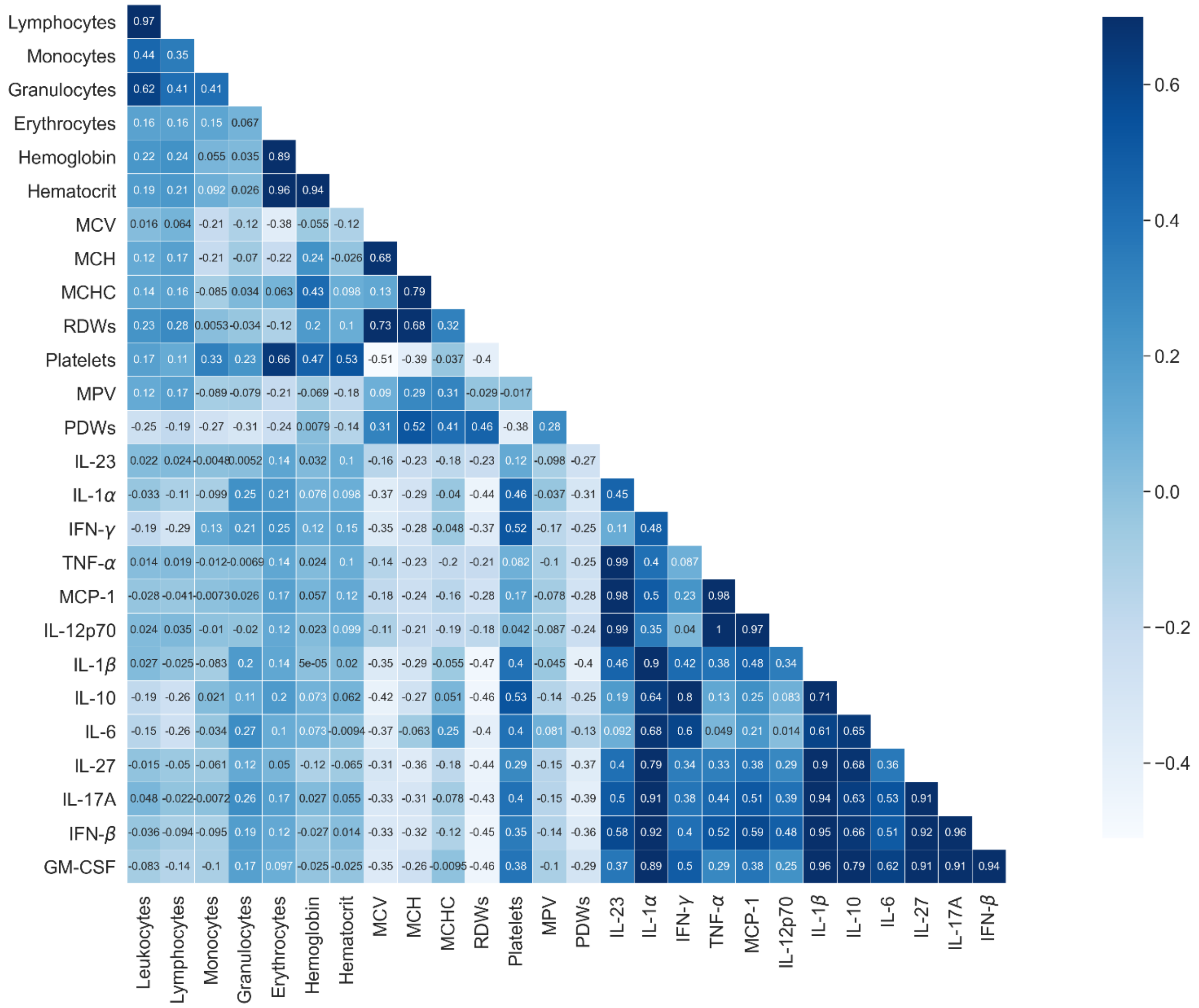


**Figure S12.** Correlations of blood cells and pulmonary cytokines over the course of influenza infection. The matrix depicts Pearson correlation coefficients of all measured blood cells and lung cytokines. Data (derived from the second experiment) were used as training data for the machine learning models.

### 3 SUPPLEMENTAL MACHINE LEARNING

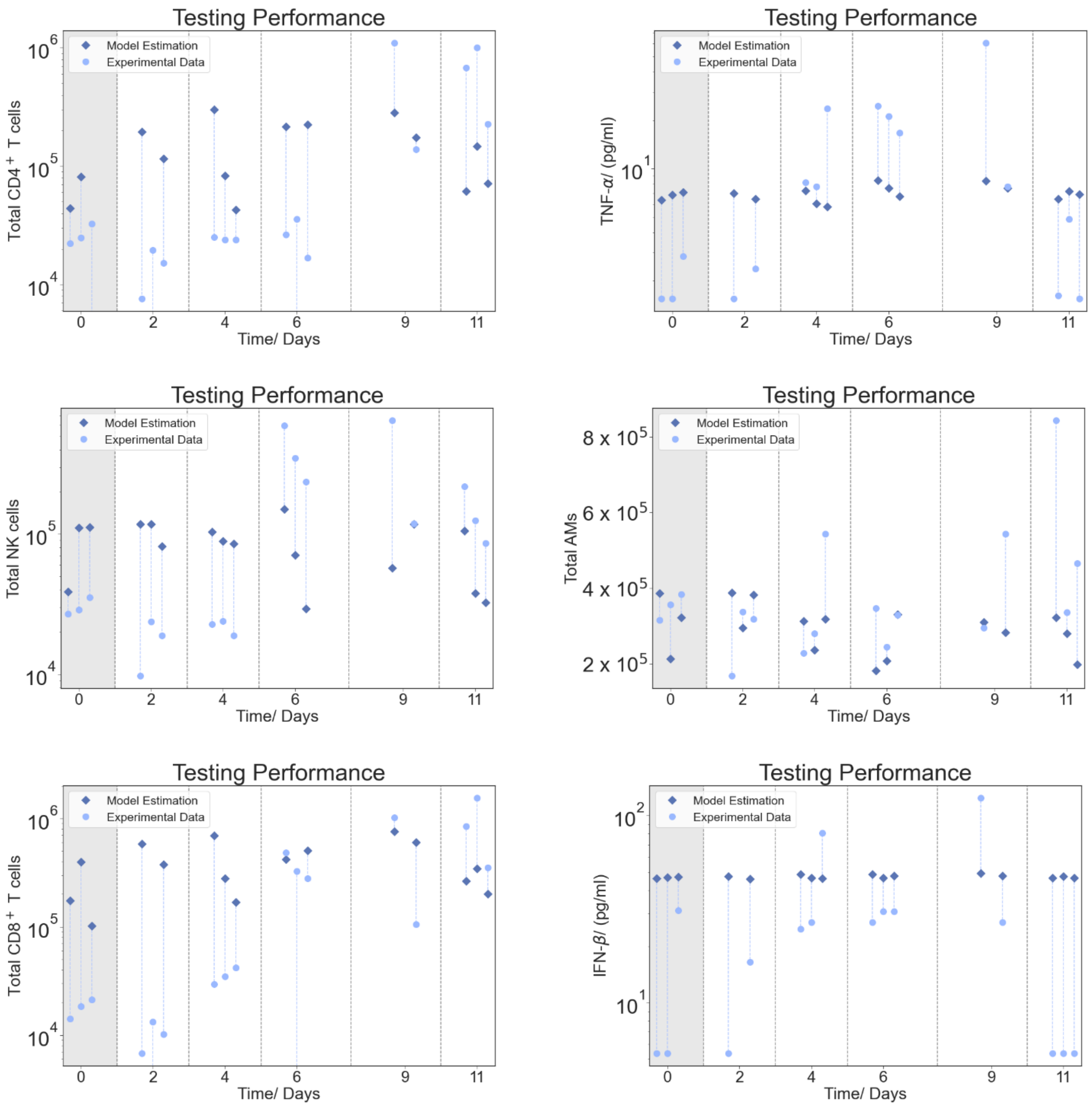


**Figure S13.** Mapping of lung leukocytes and cytokines from hematological data. Circles show the experimental data and squares the model prediction. Day 0 represents the control group. Different machine learning methods ranging from linear regression to support vector machines were applied. Here, the prediction of the best performing model is plotted, respectively.

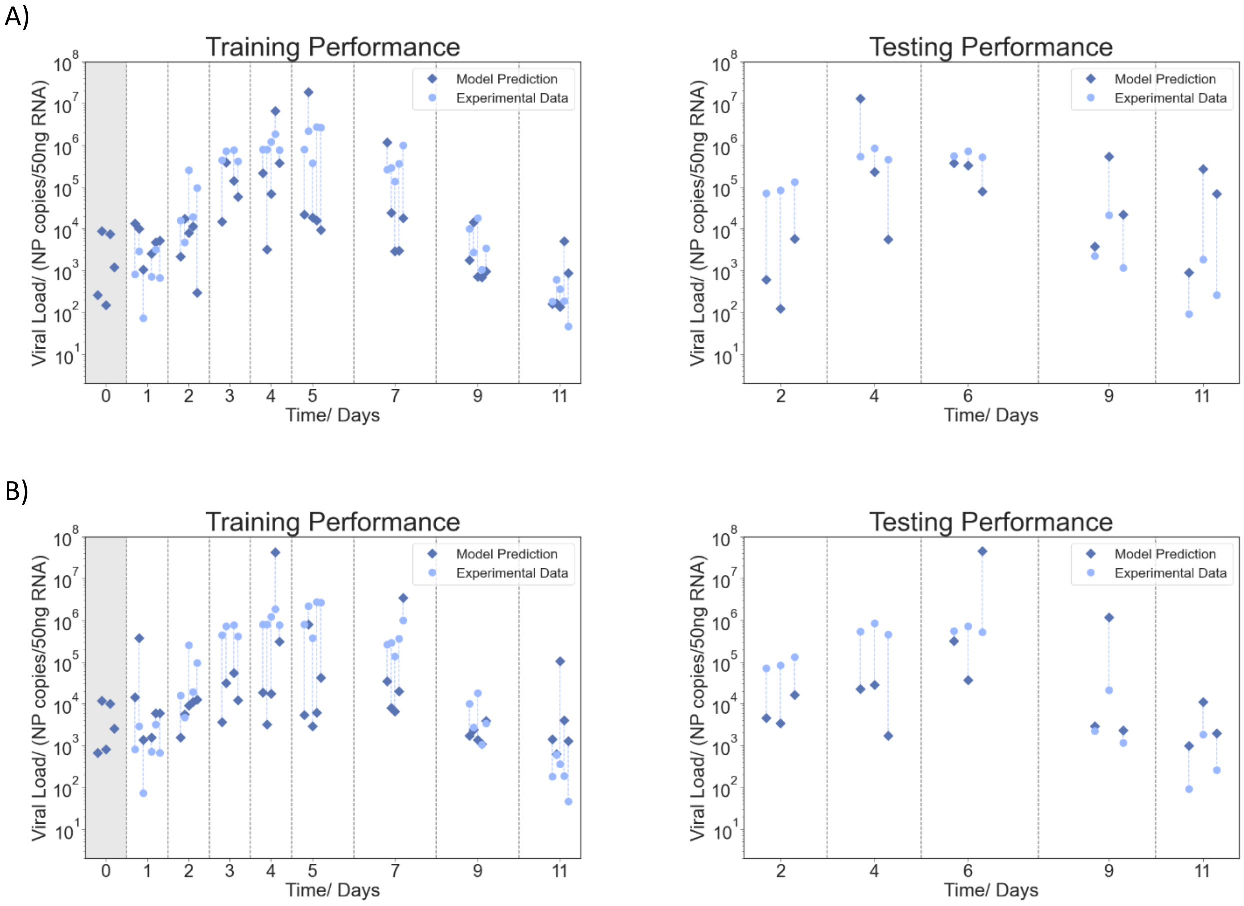


**Figure S14.** Mapping of viral load from hematological data using different features. Circles show the experimental data and squares the model prediction. Day 0 represents the control group. A) shows when only granulocytes and lymphocytes are used as input for the neural network model and all other parameters are dropped. B) shows when only the granulocyte-lymphocyte ratio and the platelet-granulocyte ratio are used as input for the neural network.

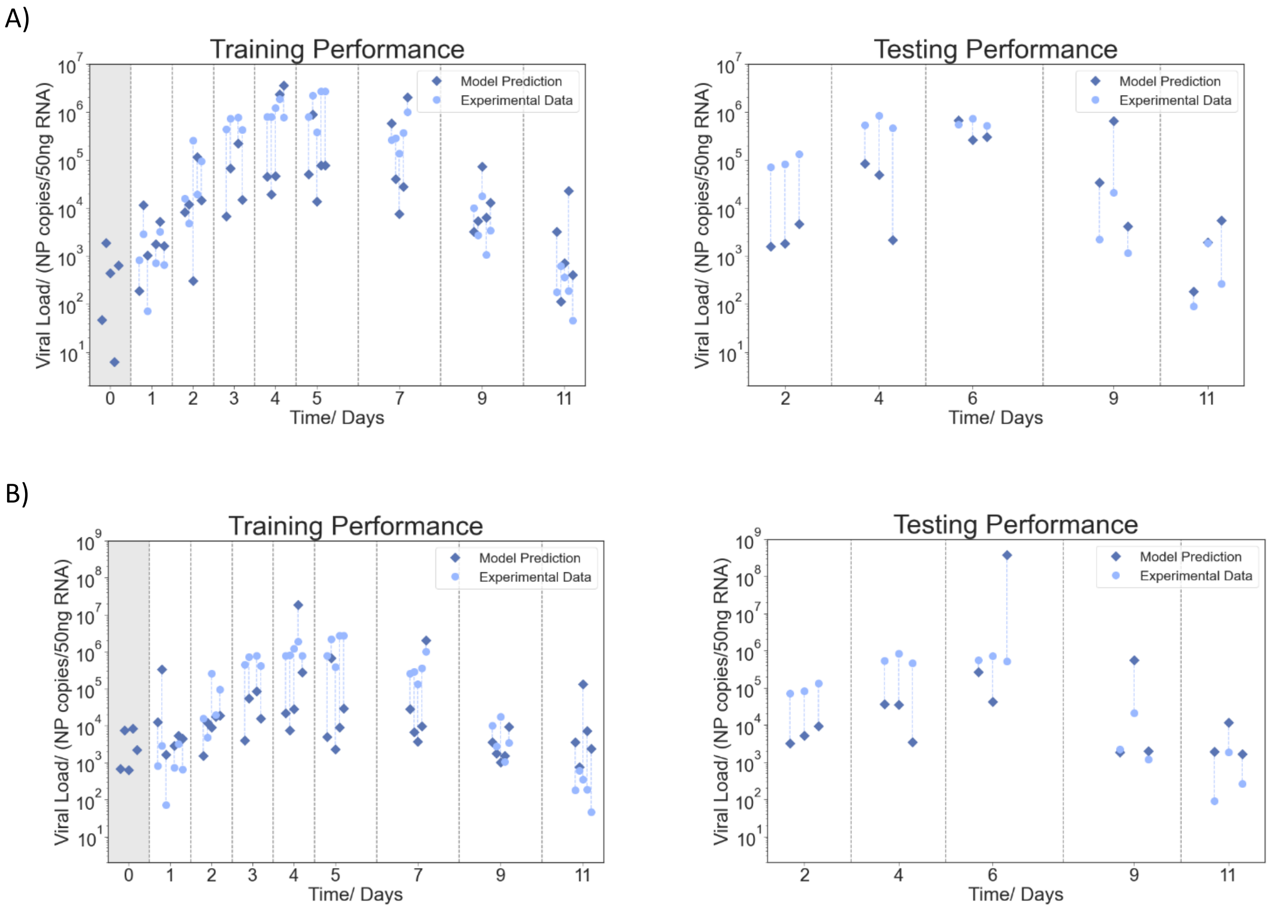


**Figure S15.** Mapping of viral load from hematological data using different features. Circles show the experimental data and squares the model prediction. Day 0 represents the control group. A) shows when granulocytes and lymphocytes are removed as input for the neural network model and replaced by the ratio of granulocytes and lymphocytes (GLR). All other parameters are unchanged, so the model has 13 features. B) shows erythrocytes, hemoglobin, GLR and platelet-granulocyte ratio (PGR) are used as input for the neural network.

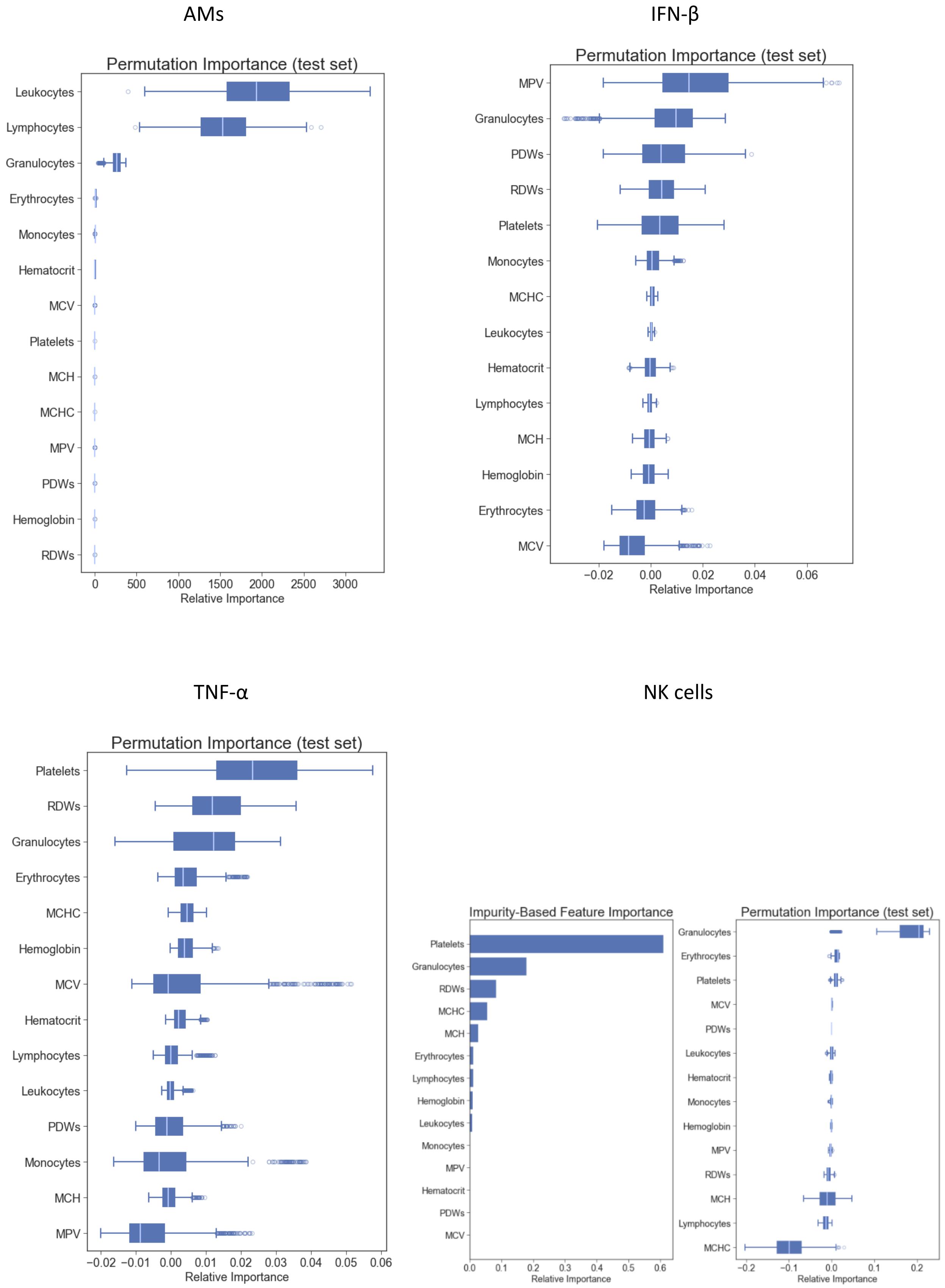


**Figure S16.** Feature importance analysis for selected lung leukocytes and cytokines from blood data. Where applicable permutation importance analysis was performed. For NK cells an impurity-based permutation analysis was additionally performed.

### 4 SUPPLEMENTAL TABLES

**Table S1**. Best performing model and respective scores for different targets from the lung milieu. Hematological data was used in all cases as input. For each variable to estimate, different machine learning models were tested. For comparison, also the $R^2$ score of the benchmark is shown. The mean of the training data was used as a benchmark.

| Target Variable | Model Type | $R^2$ | $R^2$ (benchmark) |
|---|---|---|---|
| $CD4^+$ T cells | LR with PCA | 0.01 | -0.01 |
| TNF-α | Support Vector Machine | -0.05 | -0.18 |
| $CD8^+$ T cells | LR with PCA | -0.13 | -0.05 |
| NK cells | GBRT | -0.13 | -0.20 |
| IFN-β | Support Vector Machine | -0.39 | -3.45 |
| AMs | LR | -0.49 | -0.65 |

**Table S2**. Comparison of different parameters as feature inputs for the viral load neural network model.

| Input Features | MSE | $R^2$ | $AIC_c$ (Training) |
|---|---|---|---|
| 14 parameters | 7.53 | 0.18 | -0.22 ± 3.98 |
| Replace Lymphocytes, Granulocytes with GLR | 7.33 | 0.20 | 5.81 ± 5.51 |
| Only Lymphocytes, Granulocytes | 13.11 | -0.42 | 18.37 ± 2.89 |
| Only GLR, PGR | 8.93 | 0.03 | 21.38 ± 2.11 |
| Only Erythorcytes, Hemoglobin, GLR, PGR | 9.00 | 0.03 | 19.68 ± 3.63 |
